# Germline hypomethylation shapes dynamic CpG reservoirs in ape genomes

**DOI:** 10.64898/2026.06.15.732472

**Authors:** Dongmin R. Son, Eddie Y.-H. Loh, Hyeonsoo Jeong, Jian Ma, Evan E. Eichler, Soojin V. Yi

**Affiliations:** Department of Ecology, Evolution and Marine Biology, University of California Santa Barbara; Santa Barbara, CA, USA; Neuroscience Research Institute, University of California Santa Barbara; Santa Barbara, CA, USA; Department of Life Sciences, Gwangju Institute of Science and Technology; Gwangju, South Korea; Ray and Stephanie Lane Computational Biology Department, School of Computer Science, Carnegie Mellon University; Pittsburgh, PA 15213, USA; Department of Genome Sciences, University of Washington School of Medicine; Seattle, WA 98195, USA; Howard Hughes Medical Institute, University of Washington; Seattle, WA 98195, USA; Department of Molecular, Cellular and Developmental Biology, University of California Santa Barbara; Santa Barbara, CA, USA

## Abstract

CpG dinucleotides are essential substrates for DNA methylation and genome regulation, yet paradoxically, methylated CpGs are lost through deamination-associated mutations. How CpG-rich sequences persist and are replenished during evolution remains unresolved, as previous sequencing technologies left structurally complex regions inaccessible. Here, leveraging complete telomere-to-telomere ape genomes, we show that peri/centromeric satellites are reservoirs of hypomethylated CpGs in sperm, and likely oocytes. This peri/centromeric hypomethylation extends into adjacent euchromatin, revealing novel centromeric hypomethylation extension domains enriched for segmental duplications, recently duplicated genes, and testis- and brain-biased expression. More broadly, young structural variants are CpG-rich and germline-hypomethylated, whereas older sequences are methylated and CpG-depleted. These findings link rapidly evolving centromeric regions to distinctive germline epigenetic state, CpG evolution, and structural and transcriptomic evolutionary innovation.

## Main Text

CpG dinucleotides are central to regulation of gene expression, development, and chromatin organization (*1–4*). Yet, paradoxically, methylated CpGs are inherently unstable, undergoing rapid evolutionary erosion via spontaneous deamination-associated mutations (*5–8*). Consequently, DNA methylation creates persistent evolutionary pressure toward CpG loss. Although this process explains CpG depletion, considerably less is known about the other side of CpG evolution: where new CpG-rich sequences arise, what determines their initial methylation state, and how newly introduced CpGs persist long enough to contribute to genome regulation and evolution (*9–11*).

Addressing these questions requires access to some of the most rapidly evolving regions of the genome, which undergo extensive sequence turnover, duplication, and lineage-specific expansion but have historically been inaccessible to comprehensive evolutionary and epigenetic analysis. Complete telomere-to-telomere (T2T) assemblies now resolve these regions (e.g., (*12–16*)), enabling direct examination of structurally dynamic genomic sequences and how DNA methylation influences the evolutionary fate of their CpGs.

In particular, pericentromeric and centromeric regions (hereafter, “peri/centromeric regions”) are among the most evolutionarily labile genomic compartments (*16–18*). Enriched in satellite repeats and segmental duplications, these regions undergo rapid sequence turnover (*14*, *19*, *20*), yet retain essential roles in chromosome segregation and the establishment of specialized chromatin domains (*21*, *22*). This contradiction is often referred to as the centromere paradox, whereby conserved chromosomal functions are maintained despite rapid evolution of the underlying DNA sequence (*17*, *19*). Accordingly, centromere identity and function depend on epigenetic mechanisms, including deposition of the histone variant CENP-A (*23, 24*) and DNA methylation (*25*, *26*). Yet, the tissue-specific landscapes and evolutionary consequences of DNA methylation across these rapidly evolving regions remain poorly understood.

Here, integrating complete T2T genomes of humans and other apes (*15*, *16*) with germline and somatic methylomes, we uncover a distinctive epigenetic feature with broad evolutionary implications: peri/centromeric satellite sequences are strikingly hypomethylated in sperm (and oocytes) relative to somatic tissues, making these regions disproportionately large reservoirs of CpGs. Moreover, this hypomethylation extends into adjacent non-satellite DNA, forming <u>c</u>entromeric <u>h</u>ypomethylation <u>e</u>xtension <u>d</u>omains (CHEDs) enriched for segmental duplications and recently evolved genes. Genes within CHEDs show pronounced expression biases toward testis and brain, traits often associated with human evolutionary innovation. More broadly, this pattern of sperm hypomethylation and preservation of newly introduced CpGs extends to recently emerged genomic sequences arising through dynamic structural evolution.

Together, these findings link structural genome evolution with germline epigenetic processes and identify rapidly evolving genomic regions as reservoirs of CpG-rich sequences, while revealing CHEDs as potential genomic environments for evolutionary innovation.

### Complete T2T ape genomes reveal germline methylation-driven CpG erosion

T2T assemblies of ape genomes (*15*, *16*) substantially expand the accessible CpG landscape, particularly within repetitive regions that were absent or unresolved in previous reference assemblies (**Fig. 1A**; **table S1).** For example, the human T2T-CHM13 v2 genome contains 2,916,746 (9.42%) additional CpG dinucleotides (CpGs) relative to hg38. The gorilla mGorGor1v2.0 assembly contains nearly 12 million additional CpGs (37.48%) compared with the previous genome assembly, including nearly 9 million in newly annotated satellites. Nevertheless, CpGs were depleted relative to expectation across ape genomes, with CpG observed/expected ratios (CpG O/E) centered around 0.2, whereas GpCs, an unmethylated dinucleotide control, were observed at expected frequencies (**Fig. 1, B; fig. S1A; table S2**). Comparisons of orthologous CpGs in apes clearly reveal rapid CpG loss at a rate approximately fivefold higher than that of GpC loss (**fig. S1B**), driven predominantly by C-to-T and complementary G-to-A transitions (**fig. S1C, table S3**). Interestingly, lineage-specifically expanded satellites, such as those associated with subterminal caps, including pCht in chimpanzees, gorillas, and bonobos, and alpha satellite in siamang (*16*) (example shown in **Fig. 1C**), form distinct CpG-rich peaks (**fig. S1A**). Therefore, while confirming widespread methylation-associated loss of CpGs, complete assemblies also demonstrate lineage-specific addition of CpG-rich sequences in rapidly evolving satellite regions.

**Fig. 1.**
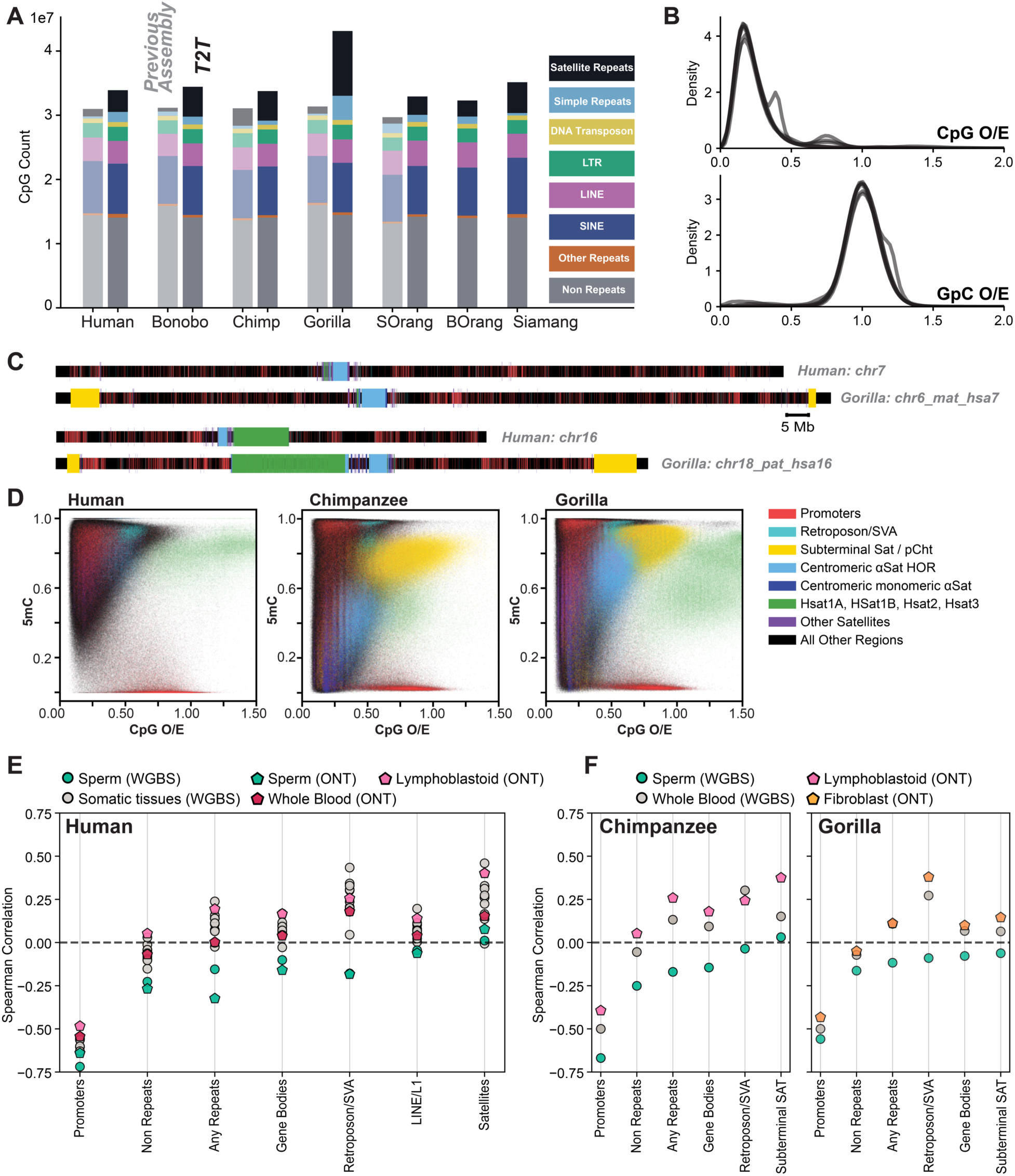
Assessment of CpG sites and DNA methylation-driven CpG evolution in ape Genomes. **(A)** Total numbers of CpGs in T2T ape genomes compared with previous assemblies, when available. CpGs are annotated by genomic context. T 5 he increase in CpG content in T2T assemblies is largely attributable to satellites and repeats. Chimp, chimpanzee; SOrang, Sumatran orangutan; BOrang, Bornean orangutan. **(B)** Distributions of CpG and GpC observed/expected (O/E) ratios in 2-kb windows across the complete genomes of seven ape species. CpG O/E shows genome-wide depletion, whereas GpC O/E is centered around 1, indicating no comparable depletion. Lineage-specific peaks in CpG O/E reflect lineage-specific satellite sequences, as shown in fig. S1. **(C)** Ideograms of genomic features across representative human and gorilla chromosomes. Satellite-related regions are shown with thicker bars. Subterminal satellites, indicated in yellow, are distinctly e 5 xpanded in gorilla compared with human. **(D)** Relationship between CpG O/E and DNA methylation across ape genomes, with representative species shown. T2T assemblies were divided into 500-bp windows, and CpG O/E and DNA methylation levels were calculated for each window. Windows were annotated by genomic feature and color-coded accordingly. High-coverage long-read DNA methylation profiles were from lymphoblastoid cells for human and chimpanzee and fibroblasts for gorilla. **(E)** Spearman correlations between CpG O/E and DNA methylation across genomic features and tissues. Highly covered windows were defined as those containing at least three CpGs, with more than 66% of CpGs covered in the corresponding tissue. Sperm consistently shows the strongest negative correlation, indicating that regions highly methylated in the germline tend to be more CpG-depleted, whereas somatic tissues generally show positive correlations. **(F)** Spearman correlations between CpG O/E and DNA methylation in chimpanzee and gorilla show a similar tissue-dependent pattern.

Different genomic features form distinct clusters of CpG O/E and DNA methylation in T2T assemblies (**Fig. 1, C and D; fig. S1D**). For example, peri/centromeric satellites are highly methylated (*27*) while relatively CpG rich in somatic-cell derived T2T assemblies (**Fig. 1D**). Because evolutionary CpG loss occurs through germline mutations, we integrated T2T assemblies with germline DNA methylation profiles as well as methylomes of primary tissues. To do so, we re-mapped a curated set of whole-genome bisulfite sequencing (WGBS) data from diverse human primary tissues of distinct developmental origins (*28–30*) and long-read methylation data from sperm and blood (*31*) to the CHM13 assembly (**table S4**). After stringent coverage and mappability filtering (Methods), 70%–80% of CpG sites were confidently covered across the WGBS tissue samples, including a subset of CpGs in highly repetitive satellite sequences (**table S5**).

Across genomic features, sperm methylation consistently shows the lowest and predominantly negative correlations with CpG O/E ratios, indicating that regions with high sperm methylation are more CpG-depleted (**Fig. 1E; table S6**). Somatic tissues, on the other hand, generally show weaker or often positive correlations, particularly in repetitive sequences (**Fig. 1E**). Promoters are an exception, exhibiting negative correlations across multiple tissues, consistent with their distinctive dynamics of epigenetic regulation (*32*, *33*). Aggregated single-cell oocyte methylomes (*34*), despite limited coverage, show broadly similar patterns to sperm (**fig. S2A**). Chimpanzee and gorilla sperm methylomes (*9*) likewise show stronger negative correlations with CpG O/E than did somatic methylomes (**Fig. 1F; table S7**). These comparisons reveal highly distinct germline methylomes relative to somatic methylomes, and identify germline DNA methylation as a major determinant of CpG content in ape genomes.

### Peri/centromeric satellites are conserved reservoirs of hypomethylated CpGs in sperm

Given the distinct patterns of germline and somatic DNA methylation, we asked whether specific genomic regions exhibit differential methylation between germline and somatic tissues. We discovered that peri/centromeric satellite sequences were strikingly hypomethylated in sperm relative to somatic tissues in both short- and long-read datasets (**Fig. 2A**). For example, the hsat(B5) subfamily shows an average 61% reduction in sperm methylation relative to B cells, and monomeric alpha satellites exhibit a 32% reduction (**Fig. 2, A to C; table S8**). Oocyte methylomes show the same tendency despite sparse coverage (**Fig. 2B; fig. S2B**). Notably, embryonic stem cells cluster with somatic tissues and exhibit relatively high satellite methylation, suggesting that the observed hypomethylation is a germline-specific feature that is reprogrammed or lost by the embryonic stem-cell stage (**Fig. 2A**). We observe similarly hypomethylated peri/centromeric satellites in chimpanzee and gorilla sperm **(Fig. 2D; table S8)**. Subterminal pCht satellites, which are specific to African great apes (*35*), also exhibit reduced methylation in sperm, as do the spacer sequences between pCht arrays (*35*) (**fig. S3**). In the human genome, which lacks pCht arrays, shorter subtelomeric satellite repeats also show sperm hypomethylation (**fig. S4**).

**Fig. 2.**
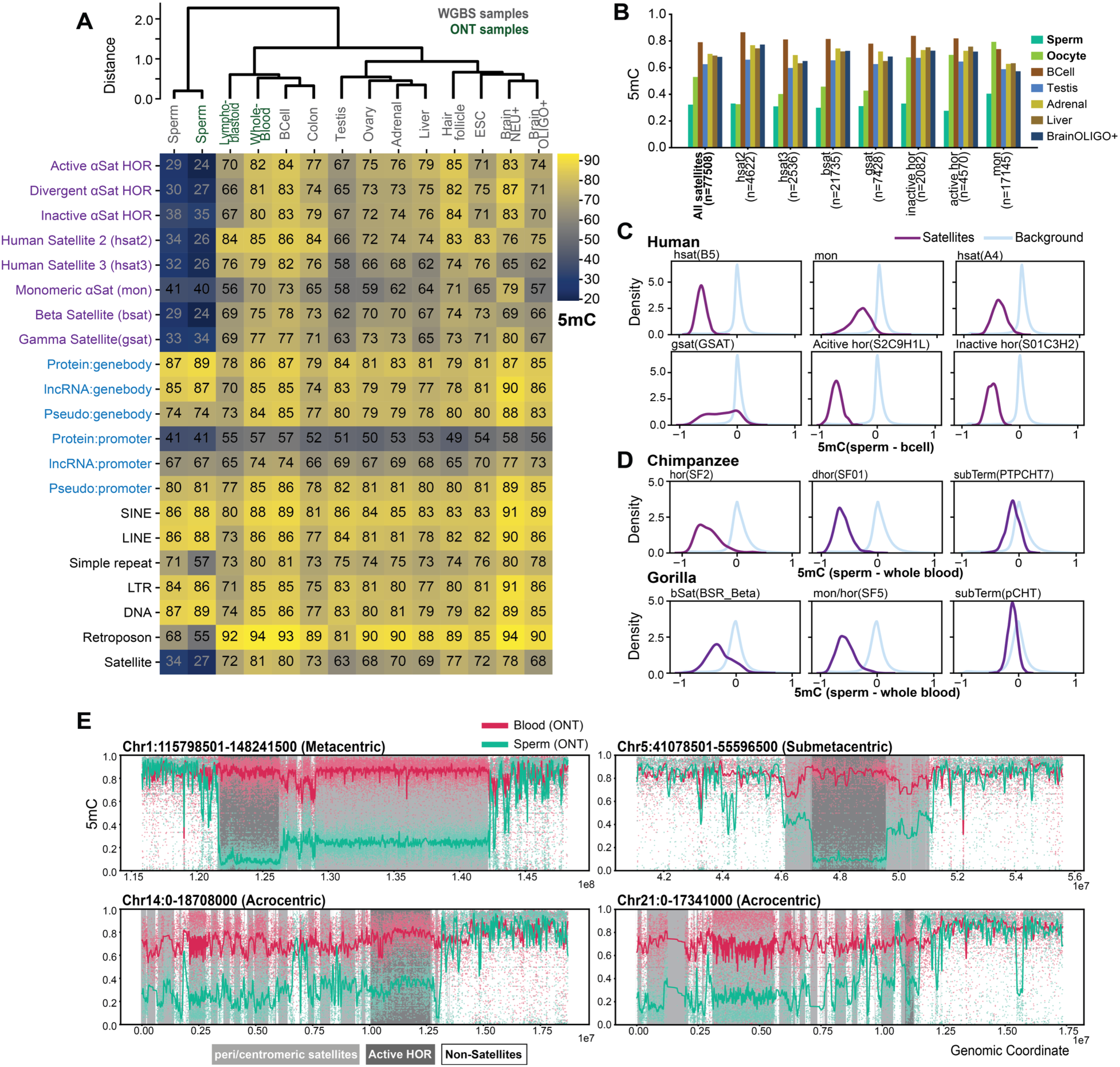
Germline-specific hypomethylation in peri/centromeric regions. **(A)** DNA methylomes across genomic contexts in diverse human tissues, hierarchically clustered by DNA methylation similarity. Sperm shows pronounced hypomethylation of peri/centromeric satellite sequences, indicated in purple. **(B)** DNA methylation of satellite sequences in sperm and oocytes compared with somatic tissues. CpGs commonly detected across all tissues, including oocytes, were compared among tissues. Numbers of CpGs analyzed are shown in parentheses. **(C)** Distribution of DNA methylation differences between sperm and B cells across representative satellite subfamilies. Satellite regions are shown in purple, and the genome-wide background is shown in light blue. **(D)** Distribution of DNA methylation differences between sperm and whole blood across representative satellite subfamilies in chimpanzee and gorilla genomes. **(E)** ONT methylation profiles of sperm (teal-green) and blood (red) across peri/centromeric regions of representative chromosomes. Gray shading indicates satellite arrays, and darker gray denotes active centromeric higher-order repeat (HOR) arrays.

Together, these results establish that peri/centromeric and subtelomeric satellites, despite their rapid sequence turnover, form a unique and ape-conserved sperm hypomethylated landscape that likely reflects a common germline pattern. Consequently, these regions harbor disproportionately higher numbers of hypomethylated CpGs in otherwise CpG-depleted ape genomes. For example, 32% of all hypomethylated CpGs in human sperm (methylation <0.3) reside within these rapidly evolving satellite sequences, even though these satellites comprise only ∼7.7% of the genome (**fig. S5**). Thus, peri/centromeric satellites constitute a disproportionately large reservoir of germline-hypomethylated CpGs.

### Satellite hypomethylation extends into adjacent non-satellite DNA

Moreover, sperm hypomethylation is not confined to satellite arrays. Both long- and short-read profiles show that hypomethylation extends beyond satellites into neighboring non-satellite sequences, whereas blood methylation remains comparatively high across these same regions (**Fig. 2E**; **Fig. 3A; fig. S6**). To delineate the extent of these domains, we compared long-read sperm and blood methylomes in overlapping 10-kb bins along each chromosome arm, extending outward from satellite boundaries until two consecutive bins no longer show significant sperm hypomethylation (Benjamini–Hochberg-adjusted FDR < 0.05). We identified approximately 37.6 Mb of hypomethylated satellite-adjacent non-satellite sequence across the human genome (**Fig.** 3B; fig. S7; table S9).

**Fig. 3.**
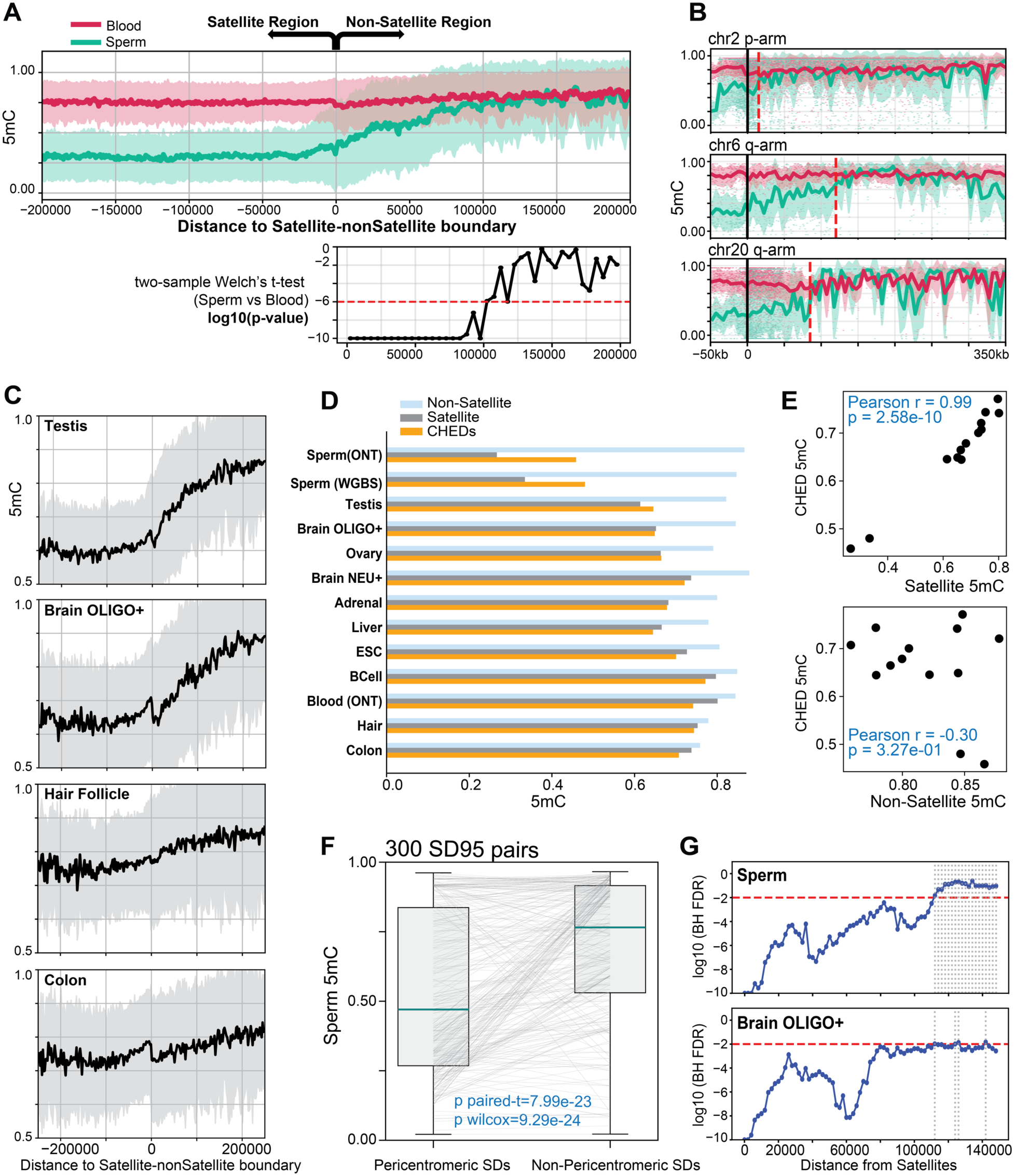
Extension of centromeric hypomethylation into adjacent genomic regions. **(A)** Sperm hypomethylation relative to blood extends beyond satellite boundaries into adjacent non-satellite sequences. Negative distances indicate satellite regions and positive distances indicate flanking non-satellite regions. Welch’s two-sample t-test significance values are shown. **(B)** Chromosome-arm specific boundaries of significant hypomethylation (red lines) were identified using overlapping 10-kb windows extending outward from satellites until two consecutive windows failed to reach significance by a one-sided Welch’s two-sample t-test with Benjamini–Hochberg correction (FDR < 0.05). Example chromosomes are shown. **(C)** DNA methylation profiles near satellite boundaries in representative tissues. Testis and oligodendrocytes (brain OLIGO+) show extended hypomethylation, whereas colon and hair follicle samples show weaker patterns. **(D)** Mean DNA methylation levels across peri/centromeric satellites, CHEDs, and background non-satellite regions across tissues. **(E)** Across tissues, CHED methylation strongly correlates with satellite methylation but not with background non-satellite methylation. Each dot represents a tissue shown in (D). Pearson correlation coefficients and P-values are shown. **(F)** DNA methylation of highly similar segmental-duplication pairs (identity >95%) with one copy near a peri/centromeric satellite boundary (0 < distance < 100 kb) and the other distal (>250 kb). Satellite-proximal copies are significantly hypomethylated in sperm. **(G)** DNA methylation divergence between SD pairs was greater when one copy was proximal to satellite regions. A sliding-window scan across 0–150 kb from satellite boundaries tested whether pericentromeric SD copies were hypomethylated relative to their corresponding distal copies. Significance was assessed using Wilcoxon signed-rank tests with Benjamini–Hochberg FDR correction, and gray dashed lines indicate FDR > 0.01.

Several tissues, notably testis and oligodendrocytes, also exhibit extended hypomethylation proximal to peri/centromeres, although to a lesser extent than in sperm (**Fig. 3, C and D; fig. S8**). In contrast, B cells, colon, and hair follicle show little to no evidence of such extended hypomethylation (**Fig. 3D; fig. S8**). To assess the consistency of these regions, we compared sperm methylation profiles to those from tissues lacking extended hypomethylation and found that the resulting domains were highly reproducible across these comparisons, with 97.67%, 94.17%, and 94.52% overlap in sperm versus B cell, colon, and hair follicle comparisons, respectively (**fig. S9)**. We refer to this reproducible, overlapping set of satellite-adjacent non-satellite regions (35.03 Mb) as centromeric hypomethylation extension domains (CHEDs) (**fig. S10, table S9**). Across tissues, CHED methylation closely tracked satellite methylation but not methylation across the remaining non-satellite genome (**Fig. 3E**).

We tested whether these domains could be explained by intrinsic sequence features rather than their proximity to peri/centromeric regions using highly similar segmental duplication (SD) pairs (>95% identity, from (*13*)) as sequence-matched controls. We show that in 300 high-confidence SD pairs (>80% of CpGs covered for both copies) in which one copy lies near peri/centromeric satellites (0 < distance < 100 kb) and the other is distal (distance > 250 kb), SD copies adjacent to peri/centromeric satellites are significantly hypomethylated relative to their distal counterparts in sperm (**Fig. 3F**). Moreover, methylation divergence between SD copies increases with proximity to satellite sequences, further supporting a distance-dependent effect (**Fig. 3G**).

Similar but weaker patterns are also observed in select somatic tissues (**figs. S11 and S12**).

Therefore, peri/centromeric hypomethylation extends beyond satellite DNA into adjacent euchromatic sequences, forming tissue-variable but reproducible CHEDs that mark dynamic epigenetic transition regions between satellite-rich heterochromatin and flanking euchromatic sequences.

### CHEDs coincide with recent structural and transcriptional innovation

Because CHEDs mark transition zones between rapidly evolving satellite arrays and flanking euchromatin, we next asked whether they also coincide with regions of structural and functional innovation. We find that CHEDs show strong associations with structural variation, gene-copy evolution, and distinctive transcriptional biases.

The extent of CHEDs was positively correlated with the abundance of SDs across chromosome arms (**figs. S10 and S17**; Spearman’s rho = 0.54, P = 0.00015). In addition, recently duplicated genes (1,793 protein-coding genes and pseudogenes, collectively referred to as SD98 genes (*36*)) are overrepresented within CHEDs. Specifically, 46.3% of 449 CHED-associated genes are classified as SD98 genes, representing a >6-fold enrichment compared with 7.2% of 22,165 genes in the rest of the genome (Fisher’s exact test, *P* < 10^-100^; **fig. S18A**, **table S14**). CHEDs also form localized reservoirs of germline-hypomethylated CpGs within these structurally dynamic regions, with a two-to threefold enrichment: although CHEDs comprise only ∼1.1% of the genome and ∼1.1% of all CpGs examined, they contain ∼2.4 to 3.3% of sperm-hypomethylated CpGs (**fig. S5**).

Consistent with the tissue-specific variability of CHEDs (**Fig. 3D**), genes residing in CHEDs also exhibit a distinctive transcriptional bias. We identified 1,909 genes (including 93 protein-coding genes, **table S10**) located within CHEDs, which we refer to as ‘CHED-associated genes’. Using RNA-seq data from 68 tissues in GTEx (*37*), we ranked gene expression levels across tissues and assessed the relative position of each tissue, where rank 1 indicates the highest expression in the tissue of interest and rank 68 the lowest. Strikingly, CHED-associated genes are disproportionately highly ranked in testis and brain compared to other genes (**Fig. 4, A and B; table S11,** Kolmogorov-Smirnov test and random-label permutation test, BH-adjusted P < 0.001 for both tests), indicating that they are preferentially utilized in these tissues. Enrichment in brain and testis is also validated in independent long-read transcriptomic datasets (Iso-Seq data) from multiple tissues (53 tissue samples, **table S12**), including sperm samples (**figs. S13 and S14, table S11**). Long-read RNA-seq data from GTEx (*38*), including multiple brain tissues, similarly support enrichment in brain (**fig. S15**).

**Fig. 4.**
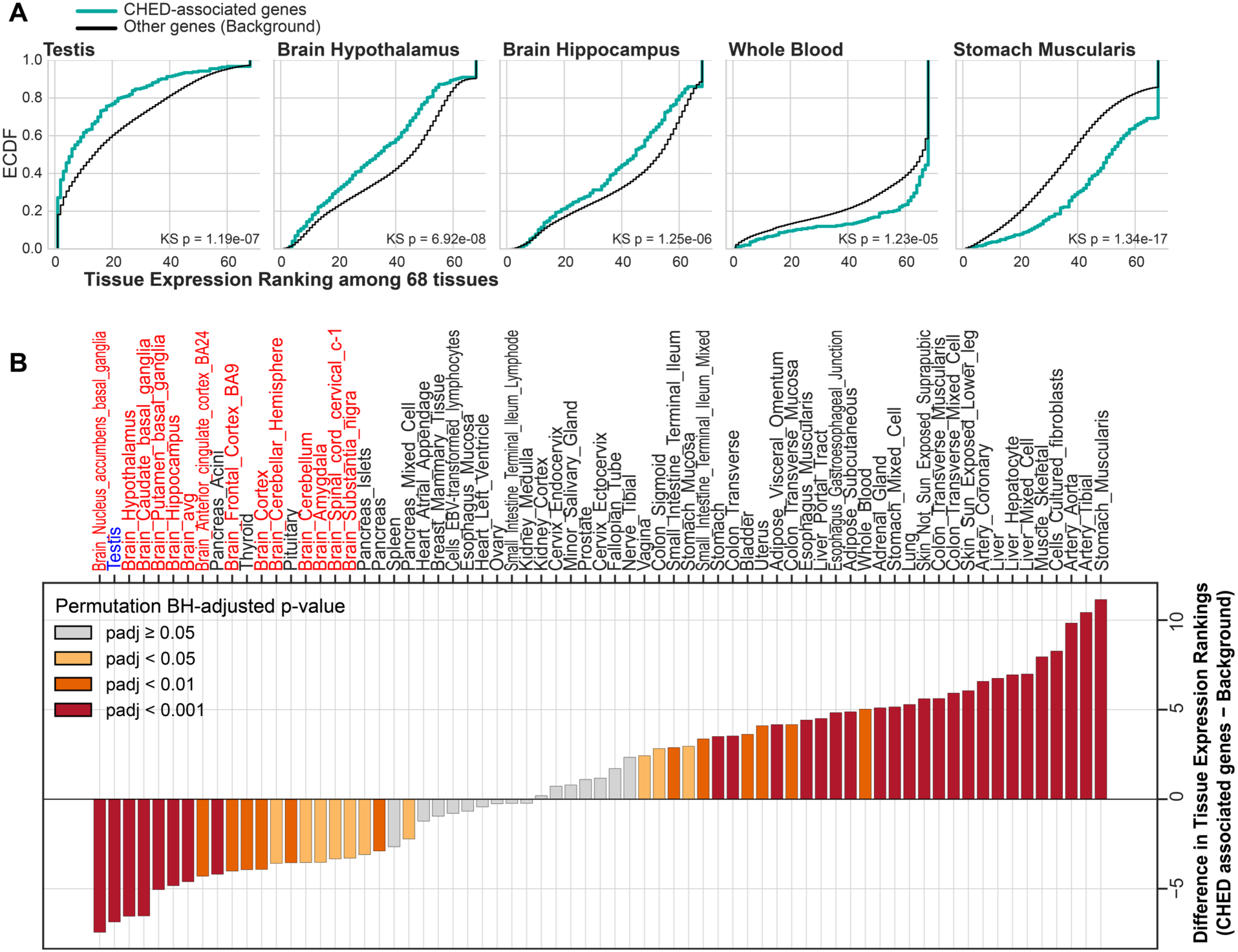
Cross-tissue expression profiles of CHED-associated genes. **(A)** Relative expression across 68 GTEx tissues was ranked from 1 to 68 for each gene, with 1 indicating the highest expression and 68 indicating the lowest. E 5 xpression-rank distributions of CHED-associated genes and background genes elsewhere in the genome were compared using ECDFs. Representative tissues are shown. In the testis and brain, CHED-associated genes showed greater enrichment than background genes, whereas other tissues, including whole blood and stomach muscularis, showed relatively low CHED-associated gene activity. P-values from Kolmogorov-Smirnov tests are indicated. **(B)** Mean expression rank differences between CHED-associated genes and background genes across 68 tissues. Negative values indicate higher relative expression of CHED-associated genes. Testis, shown in blue, and brain tissues, shown in red, exhibit distinct enrichment of CHEDassociated genes. Statistical significance was assessed using a two-sided permutation test with 10,000 random label permutations per tissue, followed by Benjamini–Hochberg FDR correction across tissues.

CHED-associated genes further exhibited coordinated tissue-biased expression and local hypomethylation. Across tissues with both methylation and expression profiles available, the degree of CHED-associated gene enrichment correlated with the extent of peri/centromeric hypomethylation (**fig. S16; table S13**; Pearson’s r = 0.56, P = 6.88 × 10⁻³), indicating that tissue-biased expression of CHED-associated genes is associated with hypomethylation of adjacent satellites. The effect of satellites is further demonstrated by an analysis of duplicate gene pairs.

Among the 84 SD98 paralog pairs in which one copy resides in a CHED and the other lies outside a CHED, CHED-associated paralogs tended to show stronger enrichment in brain and testis than their non-CHED counterparts across datasets (**fig. S18B**). CHED-associated copies also frequently exhibited reduced promoter methylation, which was associated with higher expression rank (**fig. S19**).

Together, these results indicate that CHEDs are enriched for hypomethylated CpGs, recent segmental duplications and duplicated genes, including human-specific duplicates, and that genes in these domains show reproducible expression bias toward testis and brain.

### Evolutionarily young genomic regions share a distinctive germline hypomethylation signature

The association of germline hypomethylation with rapidly evolving satellites and CHEDs suggests that evolutionarily young genomic regions may recurrently introduce CpG-rich sequences that initially experience protective germline hypomethylation. To test whether this pattern generalizes beyond satellite DNA, we examined lineage-specific segmental duplications (SDs) and simple insertions identified from T2T whole-genome multiple alignments (*13*) across the phylogeny (**Fig. 5A**; Methods).

**Fig. 5.**
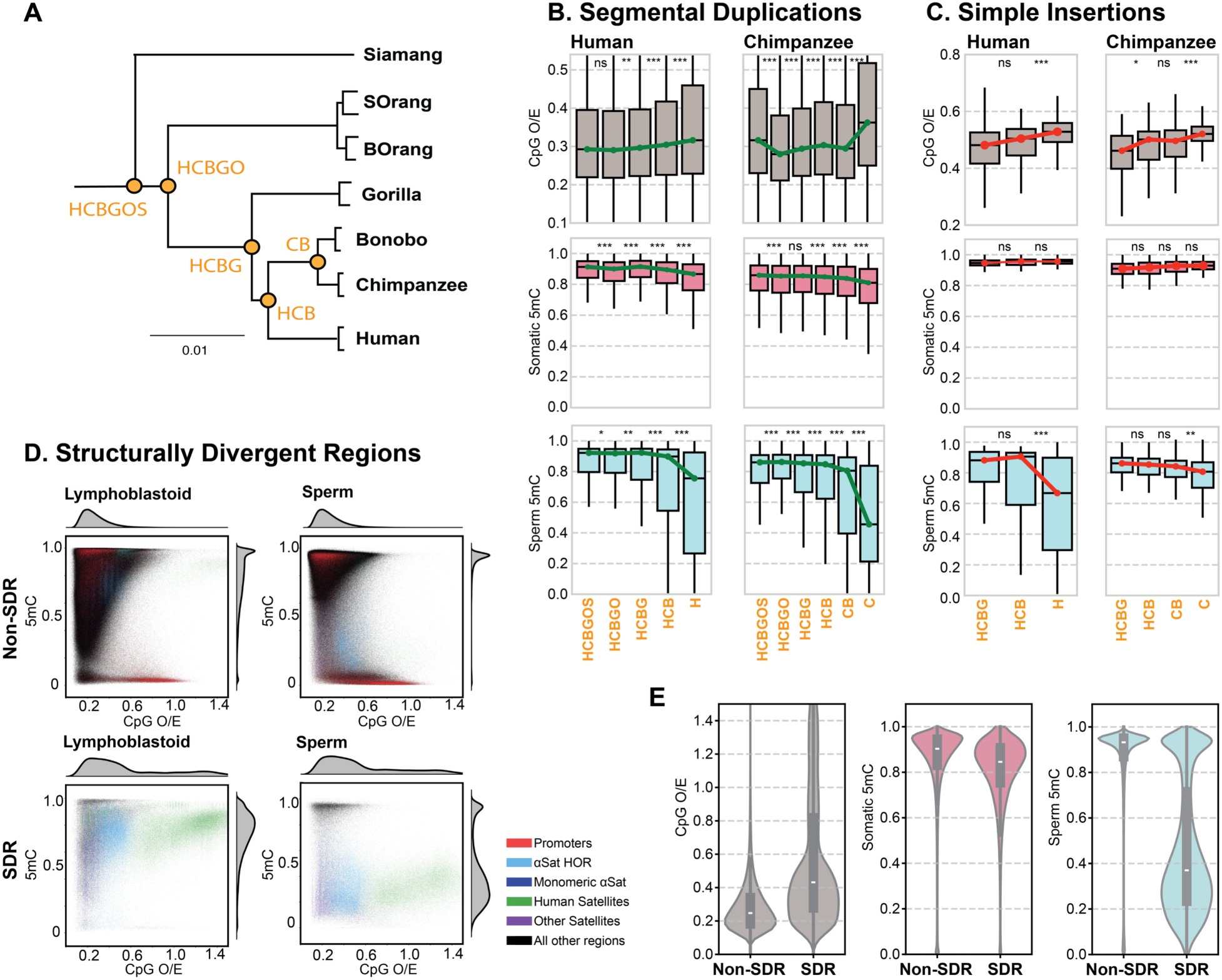
Germline hypomethylation in evolutionarily recent sequences. **(A)** Phylogenetic relationships among the species used in this study, with evolutionary nodes labeled by orange letters **(B–C)** Shared and lineage-specific segmental duplications (B) and simple insertions (C) were identified at each evolutionary node, with older elements positioned further left. CpG O/E (gray) and DNA methylation in somatic cells (red; human B cells and chimpanzee whole blood) and sperm (blue) showed higher CpG O/E and pronounced sperm hypomethylation in lineage10 specific elements. Because simple insertions are heterogeneous, sequence composition was characterized; Retroposon/SVA-associated elements are shown here. Adjacent groups were compared using two-sided Mann–Whitney U tests: *P < 0.05, **P < 0.01, ***P < 0.001; ns, p ≥ 0.05. **(D)** Actively evolving structurally divergent regions (SDRs) and non-SDRs in humans were compared for CpG O/E and DNA methylation in lymphoblastoid cells and sperm using 500-bp windows across the corresponding genomic coordinates. **(E)** CpG O/E and DNA methylation were compared between human SDRs and non-SDRs. SDRs show reduced CpG depletion (gray) and pronounced sperm hypomethylation (blue), a pattern not observed in somatic B cells (red).

We found that evolutionarily recent SDs were significantly more CpG-rich than older SDs (two-sided Mann–Whitney U test; *P* < 10^-6^ for human-specific SDs compared with those shared between humans and chimpanzees). Many lineage-specific SDs exhibit distinct hypomethylation in sperm, whereas this pattern is not as pronounced in somatic tissues (**Fig. 5B; fig. S20A**).

Evolutionarily recent simple insertions also retain higher CpG content and lower sperm DNA methylation than older, shared insertions, particularly in humans (**Fig. 5C; fig. S20B**). Because these insertions are highly heterogeneous, we further characterized and compared recurrent sequence classes within them, including SINE/Alu, LINE/L1, and retroposon/SVA elements, and observed a broadly consistent pattern across all species examined (**Fig. 5C; fig. S21; table S15**). Structurally divergent regions (SDRs), which lack clear orthologous counterparts in closely related species and are enriched for satellites and other rapidly evolving sequences (*16*), are likewise strongly hypomethylated in sperm relative to syntenically conserved regions (**Fig. 5, D and E; fig. S22)**.

Moreover, the transition regions between the extensive subterminal caps of chimpanzee and gorilla and flanking euchromatic regions, which have been shown to be enriched in recently duplicated genes (*35*), were also significantly hypomethylated in sperm relative to blood and exhibited tissue-specific variability (**fig. S23**).

Together, these results consistently show that recently emerged genomic sequences in humans and other great apes tend to be CpG-rich and relatively hypomethylated in sperm. With increasing evolutionary age, these sequences show higher sperm methylation and progressively lower CpG content. These observations link the epigenetic state of the germline to the long-term evolutionary trajectories of novel genomic sequences, suggesting that CpG birth, preservation, and erosion are coupled to the changing methylation state of structurally dynamic regions.

## Discussion

In this study, we reveal a distinctive germline methylation landscape across structurally dynamic regions of ape genomes and uncover dual roles for germline DNA methylation in shaping CpG evolution. Across most of the genome, germline DNA methylation is associated with CpG erosion, consistent with methylated CpGs being the main source of mutations via deamination of methylated cytosines (*5*, *6*, *39*, *40*). The genomic sources and early evolutionary trajectories of newly arising CpGs, on the other hand, have remained unresolved (*9–11*). Here we show that rapidly evolving peri/centromeric satellites, which were inaccessible by sequencing until recently, introduce relatively CpG-rich sequences into ape genomes. Short-read sequencing studies observed hypomethylation of centromere-proximal satellites in ape and mouse germ cells (*9*, *41*), although the lack of resolved centromeric regions precluded assessment of the extent, variability and functional consequences of this hypomethylation. Leveraging fully resolved centromeric regions of complete T2T genomes, we show that sperm hypomethylation extends broadly across most peri/centromeric satellites, including previously inaccessible centromeric sequences and active higher-order repeats (HORs), and further links this epigenetic landscape to CpG evolution and to novel methylation and transcriptional features of adjacent genomic regions. Nevertheless, it is worth noting that both female and male mouse germ cells exhibited hypomethylation near centromeres (*41*), suggesting that hypomethylation of peri/centromeric satellites may reflect a broad phylogenetic pattern.

Across complete peri/centromeric regions, this hypomethylated state creates substantial reservoirs of germline-hypomethylated CpGs. For example, nearly one third of all sperm hypomethylated CpGs in the human genome are found in these regions (**fig. S5**). Consequently, those new CpGs within these regions may be temporarily shielded from DNA methylation-driven mutations and transiently preserved. Indeed, a previous analysis of mutational profiles observed that C to T transitions were suppressed in peri/centromeric regions (*42*). In addition, satellite-rich acrocentric short arms exhibit distinctively reduced C to T transitions (*43*).

Analyses of structurally divergent regions across deeper ape phylogeny demonstrate that the relative CpG-richness and sperm hypomethylation of evolutionarily younger sequences can be generalized beyond peri/centromeric satellites, applicable to a broad range of structurally divergent genomic regions. We also show that as these sequences age evolutionarily, they acquire increased sperm methylation and subsequently undergo CpG erosion.

Remarkably, satellite hypomethylation impacts epigenetic landscapes beyond satellites themselves. In sperm, and to a lesser extent in some somatic tissues, hypomethylation extends into adjacent non-satellite euchromatic DNA to form centromeric hypomethylation extension domains, or CHEDs (**Fig. 3; fig. S8**). CHED methylation is not explained by intrinsic sequence features. Rather, it tracks nearby satellite methylation across tissues, indicating that their proximity to satellites is a main driver of their epigenetic state.

Interestingly, CHEDs, straddling heterochromatic satellite arrays and euchromatic sequences, exhibit signatures of transcriptional innovation in human evolution. They are enriched for recently duplicated genes, including human-specific duplicates. CHED-associated genes show reproducible enrichment of expression in testis and brain across different datasets. These findings extend earlier observations that centromeric transition regions and subtelomeric regions preferentially house segmental and gene duplications, and exhibit expression bias in testis (*44*–*46*). Our study offers a mechanistic context for these observations by uncovering a unique germline epigenetic profile that may prove permissive for duplicated genes and tissue-biased expression.

The enrichment of CHED-associated gene expression in testis is consistent with the longstanding view of the testis as a permissive environment for transcriptional and evolutionary innovation (*47*, *48*). Testis is known to exhibit broad transcription, potentially driven by active chromatin remodeling, as well as transcript surveillance during spermatogenesis (*49*, *50*). We also demonstrate that human CHED-associated genes are enriched in brain expression. The enrichment of CHED-associated genes in brain is notable given the extensive regulatory and structural innovation that has accompanied human brain evolution. Notably, the human brain has undergone extensive cis-regulatory remodeling during evolution, including the gain or divergence of enhancers in cortical development and adult brain (*51–53*). These regulatory innovations include elements derived from repetitive or recently evolved sequences (*54–56*).

The mechanistic basis of germline hypomethylation at peri/centromeric and subtelomeric regions remains unresolved. Hypomethylation of CHEDs tracks the degree of peri/centromeric hypomethylation, and is most pronounced in tissues with dynamic chromatin regulation (such as sperm, testis, and brain cell lineages), while less prominent in tissues where DNA methylation is known to be stable (liver, colon, and blood cells). Hypomethylation of sperm and testis may be related to dynamic epigenetic reprogramming associated with spermatogenesis (*4*, *57*, *58*).

Although a complete epigenetic reprogramming such as observed in germ cells is not present, brain development is also marked by highly dynamic and cell-type-specific epigenetic landscapes (*59–61*) and brain-derived cell lineages are known to undergo extensive chromatin remodeling and dynamic DNA methylation reprogramming (*62*, *63*), whereas many differentiated somatic tissues maintain comparatively stable, cell-type-specific epigenomic landscapes (*2*, *64*, *65*).

Together, our findings establish germline DNA methylation as both an erosive and preservative force in ape genome evolution. While driving long-term CpG erosion, reduced germline DNA methylation defines a distinctive epigenomic landscape across satellite and other structurally dynamic regions, where newly born CpGs are transiently protected from methylation-associated decay. This hypomethylated state embedded within the most rapidly evolving regions extends into neighboring euchromatic sequences, forming dynamic transition regions between heterochromatin and euchromatin that harbor potential for evolutionary innovations.

## Acknowledgments

We thank the T2T Consortium for providing access to complete genome assemblies and diverse resources. We also thank Adam Phillippy, Arang Rhie, Nancy Hansen, Steven Solar, and Brandon Pickett at NHGRI/NIH, and DongAhn Yoo, for supporting basecalling of the nanopore dataset and DNA methylation tracks in the T2T ape genomes. We are grateful to Daniela C. Soto for discussions on processing the human-specific duplicated genes and to Yang Zhang and Wendy Yang for discussions on epigenetic divergence.

## Data, code, and materials availability

Genome assemblies and associated annotation tracks were obtained from the UCSC Genome Browser and T2T Consortium resources. Ape T2T cell-line DNA methylation tracks are available through the UCSC Genome Browser.

Sources for the whole-genome bisulfite sequencing, long-read methylation, RNA-seq, and Iso-Seq datasets used in this study are listed in tables S4 and S12. GTEx expression data and ENCODE DNA methylation data were obtained from their respective public repositories, as cited in the manuscript. Derived datasets, including centromeric hypomethylation extension domain (CHED) coordinates, CHED-associated gene lists, and evolutionary sequence analyses, are provided in the supplementary tables.

## Materials and Methods

### Evaluating CpG counts and annotations in ape assemblies

We collected the T2T genomes and previous genome versions from the UCSC Genome Browser (Table S1), along with the corresponding annotation tracks. We counted all CpG sites in each reference genome and annotated them based on overlaps with the RepeatMasker track retrieved from the UCSC Genome Browser for each assembly. To calculate the CpG observed/expected (O/E) ratio, we binned each genome into 2 kb non-overlapping windows and counted CG dinucleotides, cytosines (C), and guanines (G), using: O/E = (count of CG) / ((count of C × count of G) / window_length). Windows with >10% unknown bases (N), or with no C or G, were excluded. To test species-specific peaks, we excluded windows overlapping species-specific satellites in the CenSat track.

### CpG and GpC dinucleotide ortholog identification and mutation analysis

For each of the 31,641,676 CpG and 122,194,343 GpC dinucleotide positions in the human HG002v1.0.1 Paternal genome, we used halLiftOver against a 16-way Cactus alignment to extract the aligned positions in each NHP primary genome. These identified dinucleotide positions in humans, as well as the NHPs, were extended by 10 bases on each side and sequences were extracted. Alignment of each corresponding pair of human-NHP sequences was conducted and orthology was determined and called when there was a minimum of 80% identity between the sequences. This alignment also revealed any mutations on the NHP dinucleotide orthologs themselves, which was then recorded and tabulated.

### Long-read DNA methylomes of humans and apes

To compare DNA methylation levels and CpG content across the whole genome (HG002v1.0.1 paternal for human and the v2 primary haplotypes for the other species), each genome was binned into 500 bp non-overlapping windows. CpG O/E and mean DNA methylation levels were calculated by averaging fractional methylation values across CpG sites within each window.

High-coverage ultra-long-read sequencing–based DNA methylation data for apes were generated as part of Yoo et al. (*16*). Briefly, raw signals were basecalled using Guppy v6.3.8 for all species with the model “dna_r9.4.1_450bps_modbases_5mc_cg_sup_prom.cfg”. Human, chimpanzee, and siamang data were generated from lymphoblastoid cell lines, whereas bonobo, gorilla, orangutan, and Bornean orangutan data were generated from fibroblasts. Reads were mapped to the T2T apes v2.0 and human HG002 genomes using Winnowmap v2.03. Counts of 5mC-modified bases at each CpG position were generated using Modkit v0.3.1 with the options - -cpg, --combine-strands, --ignore-h (--preset traditional), and --filter-percentile 0.2, which removes 5hmC probability and filters out the 20% of modification calls with the lowest confidence. Since 5mC in the CpG context is symmetrical, the two cytosines in a CpG context were combined to represent a single CpG site. The fractional methylation level at each CpG site was computed for sites with at least five reads, defined as the ratio of total modified base counts to the sum of modified base counts, alternative modification calls, and canonical base counts.

These tracks are available on the UCSC Genome Browser (HG002v1.1 for human and v2 assemblies for the other species). Windows were annotated with genomic features using the corresponding tracks, RefSeq, RepeatMasker, and CenSat annotations in the respective assemblies (*16*). For human, the ONT-based lymphoblastoid methylation track on HG002 was lifted over to CHM13v2.0/hs1 using UCSC LiftOver, and methylation levels were averaged across both haplotypes.

### Human DNA methylation profiles of germline and somatic tissues

To study diverse human tissues, we further collected and processed whole-genome bisulfite sequencing (WGBS) datasets curated by Mendizabal and Yi (*28*), along with WGBS data from brain, testis, and oocytes (*29*, *30*, *34*). High-coverage sperm and B-cell datasets were obtained from additional sources (*66*, *67*). The placenta was excluded from further analysis due to its unique DNA methylation profile and its mixture of fetal stem cells and adult cells.

Raw sequencing reads were trimmed using Trim Galore v0.6.10 (*68*). Reads from each tissue were aligned to the CHM13v2 (hs1) reference genome and deduplicated using Bismark v0.24.2 (*69*) with default parameters. Reads with mapping quality <20 were excluded, and secondary and supplementary alignments were removed using samtools v1.16.1. CpG-level DNA methylation was summarized using bismark_methylation_extractor. CpG sites with read depth <4 and sites with extreme coverage (above the 99th percentile) were removed. We further filtered CpG sites based on mappability to exclude sites in highly repetitive regions, defined as those with a minimum unique k-mer length >200 in both the Left-Anchored and Right-Anchored tracks retrieved from the UCSC Genome Browser (*70*). The numbers of CpG sites retained after filtering are summarized and reported. For oocytes, DNA methylation levels were estimated by aggregating reads from single-cell methylomes across the GV, MI, and MII stages, because the global average CpG methylation level is stable across these stages (*34*).

To validate hypomethylation in sperm at peri/centromeric regions, given the limited performance of WGBS in these repetitive satellite regions, we also processed nanopore-based DNA methylomes from sperm and blood (*31*). Reads and methylation tags were extracted from the raw BAM files using samtools fastq -F 2304 -T MM,ML. The reads were then mapped to the hs1 genome using minimap2 v2.30 -ax map-ont -y and sorted with samtools. The output files were used to summarize CpG-level DNA methylation using modkit pileup v0.6.1 with the options --cpg --combine-mods --combine-strands --modified-bases 5mC 5hmC, making them comparable with the WGBS dataset.

### Correlation test between genome-wide CpG O/E and DNA methylation

We computed mean methylation in 500-bp windows by averaging fractional methylation values across CpG sites within each window for each tissue. Using windows with at least three CpG sites and sufficient coverage in each tissue (>66% of CpG sites covered), we assessed correlations between CpG O/E and DNA methylation across genome-wide windows, as well as within subsets of windows overlapping specific genomic features, including diverse repeat classes, using Spearman, Pearson, and Kendall correlations (with p-values). To account for potential confounders, we also performed and reported tests that included GC content and GC skew as covariates, and computed partial correlations controlling for these variables.

For comparisons involving oocytes, because the oocyte scBS-seq data had relatively low coverage, we restricted cross-tissue correlation analyses to windows with sufficient oocyte coverage, defined as ≥3 CpG sites per window and nonzero read depth for >66% of CpGs. For chimpanzee and gorilla, we also collected WGBS datasets from whole blood and sperm and applied the same processing and analytical workflow.

### Centromeric hypomethylation in the germline

Across tissues, we compared DNA methylation levels across diverse genomic features using 500 bp windows annotated with corresponding tracks from the UCSC Genome Browser, RefSeq, RepeatMasker, and CenSat annotations on hs1 (*14*). Chromosomes X, Y, and M were excluded from the comparison.

To assess the degree of hypomethylation in sperm, we compared sperm with B cells, which showed no hypomethylation in satellite regions. Using windows detected in both tissues (≥3 CpG sites per window and >66% of CpG sites covered in both tissues), we compared and reported DNA methylation levels for satellite subfamilies represented by more than 50 windows (≥25,000 bp). We report the fraction of detected windows relative to all windows for each satellite subfamily.

To test whether detected windows were biased toward specific sequences within satellite regions, we also report differences in GC content and CpG density between detected windows and all windows. The sperm hypomethylation summary was also generated by comparing ONT-based sperm with ONT-based whole blood. For chromosome-wide visualization, we summarized methylation in 500 bp genomic windows and overlaid a Gaussian-smoothed trend line computed from the windowed values (Gaussian kernel, σ = 20 windows). Same comparisons were performed for chimpanzee and gorilla by comparing sperm and whole-blood methylation profiles, including subterminal satellite regions.

### Extension of centromeric hypomethylation into adjacent genomic regions

We first compiled a list of satellite–non-satellite boundaries at transitions between satellite arrays and non-satellite regions. Satellite arrays in contact with each other were merged into groups, and arrays shorter than 2 kb were filtered out to reduce noise from small arrays. We then collected 500 bp windows across the genome and calculated their distance from the nearest satellite–non-satellite boundary. Larger absolute values indicate greater distances from the boundary, with negative values denoting satellite regions and positive values denoting non-satellite regions. To test the significance of sperm hypomethylation extending into nearby non-satellite regions, we used sliding 5 kb bins to compare ONT-based sperm and blood methylation levels using an independent two-sample Welch’s t-test.

To further estimate the extent of hypomethylation extending into nearby non-satellite regions on each chromosome, we collected 500bp non-satellite windows and calculated the distance from each window to the nearest satellite–non-satellite boundary. Autosomal chromosomes were separated into p and q arms based on centromere position, defined as the midpoint of the active HOR array on each chromosome. This allowed us to compare DNA methylation patterns in nearby non-satellite regions as a function of their distance from satellite sequences.

Using ONT-based sperm and blood methylation data, we estimated the distance over which sperm hypomethylation remained statistically significant relative to blood using a sliding-bin approach. We first collected non-satellite windows located within 0–10 kb of satellite sequences and tested for sperm hypomethylation using a one-sided Welch’s two-sample t-test (sperm < blood), using the window mean methylation values in the range. We then repeated this test in sliding 10 kb bins with 5 kb overlaps, including 0–10 kb, 5–15 kb, 10–20 kb, and subsequent intervals extending up to 400 kb from satellite sequences for each chromosome arm. Sliding bins containing fewer than 10 data points were excluded from testing. P-values were corrected for multiple testing using the Benjamini–Hochberg false discovery rate (FDR) procedure. Significant hypomethylated range was defined as the distance interval extending from the satellite boundary until two consecutive bins were no longer significant (q ≥ 0.05). Genomic windows within this significant range were then collectively defined as CHEDs.

The same analysis was repeated using WGBS samples by comparing sperm with somatic tissues showing no or low levels of centromeric hypomethylation, including B cells, colon, and hair follicle samples. CHEDs were identified using the same sliding-bin approach for each pairwise comparison, and their overlap with ONT-based CHEDs was assessed based on genomic coordinates to evaluate consistency. CHEDs supported by all pairwise comparisons were collectively used for downstream analyses. Because sex chromosomes are known to exhibit distinct sex-specific methylation patterns across samples, CHEDs were defined only for autosomes. We also calculated average DNA methylation levels within the defined CHEDs across 13 human tissues and compared them with those in satellite and non-satellite regions to validate whether CHEDs showed methylation patterns more similar to satellite regions than to other non-satellite regions.

### Segmental duplication

For segmental duplication (SD) analyses on the human genome hs1, we used SD annotations from Vollger et al. (*13*), retrieved from the UCSC Genome Browser. We calculated the mean DNA methylation level for each SD by averaging methylation across CpG sites within the SD across tissues. We then compared highly identical SD pairs (fracMatch > 0.95) in which one copy is proximal to, but not within, peri/centromeric regions (distance to the nearest Satellite– nonSatellite boundary, 0–100 kb) and the other copy is distal (>250 kb). We further restricted analyses to SD pairs with high coverage, requiring that in the tissue examined, >80% of CpG sites were covered in both copies. For each pair, we computed the methylation difference as the methylation level of the SD copy near the satellite minus that of the SD copy distal to the satellite. We tested whether these paired differences were significantly less than zero across loci (i.e., whether the Satellite-proximal copy tends to have lower methylation), using a one-sided one-sample t-test and a one-sided Wilcoxon signed-rank test.

We also used these SD pairs to test for a distance-dependent effect, assessing whether DNA methylation divergence between SD copies depended on distance from satellite sequences. For tissues with more than 2,000 covered SD pairs overall, we performed a sliding-window scan across highly identical SD pairs with one copy near a satellite region (40-kb windows, 2-kb step size, spanning 0–150 kb from the satellite boundary). For each window, we compared pairs in which the satellite-proximal copy fell within the window and the corresponding distal copy was ≥250 kb from the nearest satellite. Windows with fewer than 10 paired observations were not tested. Within each tested window, we applied a one-sided paired Wilcoxon signed-rank test (alternative: satellite-proximal copy < distal copy) to compare mean DNA methylation levels. We then corrected p-values across all tested windows within each tissue using the Benjamini– Hochberg FDR procedure (α = 0.01). Windows were considered significant if the FDR-adjusted p-value was <0.01, and we visualized window-wise adjusted p-values on a log scale as a function of genomic distance.

### Cross-tissue gene expression profiles for CHED-associated genes

Genes in human T2T-CHM13v2.0 were obtained from the UCSC GENCODEv35 CAT/Liftoff v2 gene annotation (*71*). Promoters were defined as 2,500 bp upstream and 500 bp downstream of gene transcription start sites. We defined CHED-associated genes as genes that overlap with the defined CHEDs. These genes are reported, along with their DNA methylation levels across tissues and CpG coverage, including for all other gene bodies and promoters.

Gene expression for these genes across 68 tissues was obtained from GTEx v10 bulk-tissue RNA-seq data (GTEx_Analysis_v10_RNASeQCv2.4.2_gene_median_tpm) by matching Ensembl gene IDs. To compare relative testis expression across the 68 tissues; we ranked testis for each gene by ordering tissue TPM values in descending order using the “max” tie method; a rank of 1 indicates that testis has the highest TPM among all tissues; whereas a rank of 68 indicates that testis has the lowest. We then compared the ECDFs of testis ranks between pericentromeric CHED-associated genes and a background set of non-CHED-associated genes.

To test whether CHED-associated genes’ expression-rank distributions differed significantly from background genes, we performed a two-sided two-sample Kolmogorov– Smirnov test comparing their empirical distributions. Because CHED-associated genes tended to have lower testis ranks (i.e., be more testis-enriched) than background genes, we quantified the signed mean difference between the two distributions (mean testis rank for CHED-associated genes minus mean testis rank for nonCHED-associated genes), which is mathematically equivalent to the area between the two ECDF curves. We performed analogous analyses for all other tissues, including brain regions, and compared the degree of CHED-associated gene enrichment across tissues. We quantified differences using the signed mean rank difference between the two distributions. For these cross-tissue comparisons, we retained only genes available in GTEx and filtered out genes whose expression was not detected (TPM ≤ 0.1) in most tissues, in more than 58 of the 68 tissues.

Statistical significance was assessed for each tissue using both a two-sample Kolmogorov-Smirnov test and a two-sided permutation test with 10,000 permutations. The Kolmogorov-Smirnov test was used to compare the full expression-rank distributions between CHED-associated genes and other background genes. For the permutation test, gene-group labels, CHED-associated or background genes, were randomly shuffled while preserving the original group sizes, and the mean rank difference was recalculated to generate a null distribution. The empirical permutation p-value was calculated as the proportion of permuted absolute mean differences greater than or equal to the observed absolute mean difference. P-values from each test were adjusted across tissues using the Benjamini–Hochberg false discovery rate correction.

To test whether variation in CHED-associated gene enrichment is associated with tissue-specific differences in peri/centromeric satellite DNA methylation, we used DNA methylation profiles from tissues in our study that overlap with GTEx. We further incorporated tissue WGBS datasets from the ENCODE Project (*29*) to generate a more comprehensive set of matched methylation–expression profiles, downloading processed BED files containing CpG methylation calls. In these files, methylation measurements from the two complementary cytosines were merged, and sample-level methylation values were averaged when multiple samples were available. CpG coordinates were converted from hg38 to CHM13 using UCSC liftOver. DNA methylation levels were summarized in 500-bp genomic windows for each tissue. We retained commonly detected windows containing at least three CpG sites and with >66% of CpG sites covered in all tissues, excluding the sex chromosomes. Using windows overlapping peri/centromeric satellites (10,702 windows; 5,351,000 bp total), we estimated peri/centromeric DNA methylation by averaging methylation levels across these windows and compared it with the genomic background (non-peri/centromeric windows). We quantified peri/centromeric hypomethylation relative to background using the normalized methylation difference: (pericentromeric 5mC − background 5mC) / (pericentromeric 5mC + background 5mC).

We then mapped tissues with DNA methylation profiles to the best-matching GTEx tissue and tested whether the degree of peri/centromeric hypomethylation across tissues correlates with the degree of CHED-associated gene enrichment across the matched GTEx tissues. Correlations were assessed using both Pearson’s correlation coefficient and Spearman’s rank correlation coefficient. For the brain samples in this analysis, we aggregated GTEx brain regions by averaging tissue expression rankings across all GTEx brain tissues.

### Long-read sequencing gene expression profiles for CHED-associated genes

To validate the enrichment of CHED-associated gene expression in testis and brain samples using independent datasets, we analyzed GTEx long-read RNA-seq data (GTEx Analysis V9; 15 tissues; (*38*)) as well as compiled Iso-Seq data across diverse tissues re-mapped to the T2T reference genome (53 tissue samples). For the GTEx long-read RNA-seq data, we retrieved transcript TPMs, summed them to obtain gene-level TPMs, and integrated the results by matching Ensembl gene IDs.

PacBio Iso-Seq data specifically for brain, ovary, and testis tissues were extracted from the comprehensive dataset published by Jeong et al. (*72*). These Iso-Seq data and other tissue datasets downloaded from public databases were processed consistently. Raw reads were mapped to the human CHM13v2 reference using minimap2 v2.22-r1101 (*73*) with -ax splice:hq - -cs --eqx -y -f 1000 -p 0.8 -N 5 --secondary=yes. Alignments were then converted to BAM format (samtools view -@ ${THREADS} -b), coordinate-sorted using samtools v1.13 (*74*), and indexed with samtools index. Gene-level counts were generated across all BAM files with featureCounts v2.0.1 (*75*) using -a ${GTF} -t exon -g gene_id -s 1 --primary -L and summarized using the UCSC GENCODEv35 CAT/Liftoff v2 gene annotation. For both datasets, TPM values were computed for each sample using the median value (or the mean if there were two samples) when multiple samples were available. For these cross-tissue expression ranking comparisons, we filtered out genes that were not detected (TPM ≤ 0.1) in most tissues, in more than 39 of the 53 tissues.

### Effect of peri/centromeric hypomethylation on duplicated genes

We assessed overlap between CHED-associated genes and the list of highly identical duplicated genes (SD98 genes) from Soto et al. (*36*) on protein-coding genes and unprocessed pseudogenes. We tested whether CHED-associated genes were enriched for SD98 genes by comparing the proportion of SD98 genes among CHED-associated genes with the proportion among genes in the rest of the genome. We also annotated whether each duplicated gene was human-specific based on synteny with chimpanzees (*36*). To test DNA methylation and expression divergence between paralogous copies, we analyzed SD98 genes that are fixed or nearly fixed in humans, whose gene coordinates fall entirely within segmental duplications (consistent with whole-gene duplication), and that belong to gene families with at least two members (*36*). We then generated paralog pairs within the same gene family in which one paralog was a CHED-associated gene and the other was not, yielding 84 gene pairs, to compare divergence between duplicated genes that include a CHED-associated paralog.

To quantify expression divergence, we compared cross-tissue expression rankings and, for each paralog pair, calculated the tissue-specific ranking difference as the ranking of the CHED-associated paralog minus that of the non-CHED-associated paralog across 68 GTEx tissues in which both copies had available cross-tissue TPM values. Negative values indicate that the CHED-associated paralog is relatively more highly expressed in that tissue than the non-

CHED-associated paralog. We compared, across tissues, the proportion of pairs in which the CHED-associated paralog had a higher tissue ranking than the non-CHED-associated paralog. The same method was applied to the GTEx long-read data and the compiled Iso-Seq dataset.

To calculate DNA methylation divergence, we restricted analyses to paralog pairs in which both copies were well covered, with >66% of CpG sites in gene bodies and promoters covered in the tissue being compared. For each pair, the normalized DNA methylation difference was computed as (5mC_CHED-associated paralog − 5mC_non-CHED-associated paralog) / (5mC_CHED-associated paralog + 5mC_non-CHED-associated paralog). Finally, to test whether DNA methylation divergence was associated with expression divergence (the mean tissue-expression rank of the CHED-associated paralog minus that of the non-CHED-associated paralog), we used the most closely corresponding tissues for which both DNA methylome and transcriptome profiles were available.

### DNA methylation in evolutionarily recent sequences

To compare germline-specific patterns in evolutionarily recent regions, we focused on human, chimpanzee, and gorilla, for which somatic tissue and sperm DNA methylation profiles were available.

Lists of shared and lineage-specific segmental duplications (SDs) were generated from Yoo et al. (*16*). Human-specific SDs identified in HG002 were lifted over to the CHM13v2 genome using UCSC liftOver. Using the previously summarized 500 bp genomic windows (with tissue DNA methylation levels and CpG O/E ratios already computed), we annotated each window as overlapping either shared or lineage-specific SDs. We then collected SD-overlapping windows in each tissue and compared DNA methylation levels and CpG O/E between shared and lineage-specific SDs.

Structurally divergent regions (SDRs) represent rapidly evolving and structurally variable regions of the genome; in ape genomes, 12.5–27.3% of the genome fails to align or is inconsistent with a simple one-to-one alignment among species (*16*). Using SDR coordinates, we annotated the 500 bp windows as SDR-overlapping or non-overlapping. For SDR-annotated windows with at least three covered CpG sites in each tissue, we compared DNA methylation levels and CpG O/E between SDR and non-SDR regions.

To identify shared and lineage-specific simple insertions across the ape phylogeny, we utilized the 16-way Progressive Cactus multiple sequence alignment. To ensure high-confidence calls and avoid alignment ambiguities arising from complex structural variations, we restricted our analysis to strictly one-to-one syntenic regions. Using the HAL toolkit (*76*), we filtered out one-to-many and many-to-one mappings, extracting only mutually orthologous genomic blocks across the primate assemblies. Within these one-to-one regions, insertions and deletions were identified using halBranchMutations, which allowed us to assign simple insertions to specific evolutionary nodes (e.g., human-specific, or shared between the human-chimpanzee ancestor).

For the identified simple insertions, we calculated tissue DNA methylation levels and CpG O/E ratios. For human, insertions on the paternal haplotype of HG002 were lifted over to the CHM13v2 genome for comparison. Simple insertions longer than 100 bp and with at least three covered CpG sites in each tissue were retained for downstream analyses. Given that simple insertions can comprise highly heterogeneous sequences derived from different classes of transposable elements, we further characterized their repeat composition using species-specific RepeatMasker annotations. Repeat elements fully contained within a simple insertion interval were identified and classified by repeat type. We then compared DNA methylation levels and CpG observed-to-expected (O/E) ratios across repeat types at each evolutionary node.

## References

1. A. Bird, DNA methylation patterns and epigenetic memory. Genes & Development 16, 6–21 (2002).

2. P. A. Jones, Functions of DNA methylation: islands, start sites, gene bodies and beyond. Nat. Rev. Genet. 201, 484–492 (2012).

3. D. Schübeler, Function and information content of DNA methylation. Nature 517, 321– 326 (2015).

4. M. V. C. Greenberg, D. Bourc’his, The diverse roles of DNA methylation in mammalian development and disease. Nature Reviews Molecular Cell Biology 20, 590–607 (2019).

5. A. Bird, DNA methylation and the frequency of CpG in animal DNA. Nucleic Acids Res. 8, 1499–1504 (1980).

6. J. Sved, A. Bird, The expected equilibrium of the CpG dinucleotide in vertebrate genomes under a mutation model. Proc. Natl. Acad. Sci. USA. 85, 2653–2657 (1990).

7. N. Elango, S.-H. Kim, NISC Comparative Sequencing Program, E. Vigoda, S. V. Yi, Mutations of different molecular origins exhibit contrasting patterns of regional substitution rate variation. PLoS Comput. Biol. 4, e1000015 (2008).

8. R. Holliday, G. W. Grigg, DNA methylation and mutation. Mutat. Res. 285, 61–67 (1993).

9. A. Molaro, E. Hodges, F. Fang, Q. Song, W. ¬. McCombie, G. Hannon, A. Smith, Sperm methylation profiles reveal features of epigenetic inheritance and evolution in primates. Cell 146, 1029–1041 (2011).

10. S. V. Yi, Insights into epigenome evolution from animal and plant methylomes. 9, 3189– 3201 (2017).

11. N. Cohen, E. Kenigsberg, A. Tanay, Primate CpG Islands Are Maintained by Heterogeneous Evolutionary Regimes Involving Minimal Selection. Cell 145, 773–786 (2011).

12. S. Nurk, S. Koren, A. Rhie, M. Rautiainen, A. V. Bzikadze, A. Mikheenko, M. R. Vollger, N. Altemose, L. Uralsky, A. Gershman, S. Aganezov, S. J. Hoyt, M. Diekhans, G. A. Logsdon, M. Alonge, S. E. Antonarakis, M. Borchers, G. G. Bouffard, S. Y. Brooks, G. V. Caldas, N.-C. Chen, H. Cheng, C.-S. Chin, W. Chow, L. G. de Lima, P. C. Dishuck, R. Durbin, T. Dvorkina, I. T. Fiddes, G. Formenti, R. S. Fulton, A. Fungtammasan, E. Garrison, P. G. S. Grady, T. A. Graves-Lindsay, I. M. Hall, N. F. Hansen, G. A. Hartley, M. Haukness, K. Howe, M. W. Hunkapiller, C. Jain, M. Jain, E. D. Jarvis, P. Kerpedjiev, M. Kirsche, M. Kolmogorov, J. Korlach, M. Kremitzki, H. Li, V. V. Maduro, T. Marschall, A. M. McCartney, J. McDaniel, D. E. Miller, J. C. Mullikin, E. W. Myers, N. D. Olson, B. Paten, P. Peluso, P. A. Pevzner, D. Porubsky, T. Potapova, E. I. Rogaev, J. A. Rosenfeld, S. L. Salzberg, V. A. Schneider, F. J. Sedlazeck, K. Shafin, C. J. Shew, A. Shumate, Y. Sims, A. F. A. Smit, D. C. Soto, I. Sović, J. M. Storer, A. Streets, B. A. Sullivan, F. Thibaud-Nissen, J. Torrance, J. Wagner, B. P. Walenz, A. Wenger, J. M. D. Wood, C. Xiao, S. M. Yan, A. C. Young, S. Zarate, U. Surti, R. C. McCoy, M. Y. Dennis, I. A. Alexandrov, J. L. Gerton, R. J. O’Neill, W. Timp, J. M. Zook, M. C. Schatz, E. E. Eichler, K. H. Miga, A. M. Phillippy, The complete sequence of a human genome. Science 376, 44–53 (2022).

13. M. R. Vollger, X. Guitart, P. C. Dishuck, L. Mercuri, W. T. Harvey, A. Gershman, M. Diekhans, A. Sulovari, K. M. Munson, A. P. Lewis, K. Hoekzema, D. Porubsky, R. Li, S. Nurk, S. Koren, K. H. Miga, A. M. Phillippy, W. Timp, M. Ventura, E. E. Eichler, Segmental duplications and their variation in a complete human genome. Science 376, eabj6965 (2022).

14. N. Altemose, G. A. Logsdon, A. V. Bzikadze, P. Sidhwani, S. A. Langley, G. V. Caldas, S. J. Hoyt, L. Uralsky, F. D. Ryabov, C. J. Shew, M. E. G. Sauria, M. Borchers, A. Gershman, A. Mikheenko, V. A. Shepelev, T. Dvorkina, O. Kunyavskaya, M. R. Vollger, A. Rhie, A. M. McCartney, M. Asri, R. Lorig-Roach, K. Shafin, J. K. Lucas, S. Aganezov, D. Olson, L. G. de Lima, T. Potapova, G. A. Hartley, M. Haukness, P. Kerpedjiev, F. Gusev, K. Tigyi, S. Brooks, A. Young, S. Nurk, S. Koren, S. R. Salama, B. Paten, E. I. Rogaev, A. Streets, G. H. Karpen, A. F. Dernburg, B. A. Sullivan, A. F. Straight, T. J. Wheeler, J. L. Gerton, E. E. Eichler, A. M. Phillippy, W. Timp, M. Y. Dennis, R. J. O’Neill, J. M. Zook, M. C. Schatz, P. A. Pevzner, M. Diekhans, C. H. Langley, I. A. Alexandrov, K. H. Miga, Complete genomic and epigenetic maps of human centromeres. Science 376, eabl4178 (2022).

15. K. D. Makova, B. D. Pickett, R. S. Harris, G. A. Hartley, M. Cechova, K. Pal, S. Nurk, D. Yoo, Q. Li, P. Hebbar, B. C. McGrath, F. Antonacci, M. Aubel, A. Biddanda, M. Borchers, E. Bornberg-Bauer, G. G. Bouffard, S. Y. Brooks, L. Carbone, L. Carrel, A. Carroll, P.-C. Chang, C.-S. Chin, D. E. Cook, S. J. C. Craig, L. de Gennaro, M. Diekhans, A. Dutra, G. H. Garcia, P. G. S. Grady, R. E. Green, D. Haddad, P. Hallast, W. T. Harvey, G. Hickey, D. A. Hillis, S. J. Hoyt, H. Jeong, K. Kamali, S. L. K. Pond, T. M. LaPolice, C. Lee, A. P. Lewis, Y.-H. E. Loh, P. Masterson, K. M. McGarvey, R. C. McCoy, P. Medvedev, K. H. Miga, K. M. Munson, E. Pak, B. Paten, B. J. Pinto, T. Potapova, A. Rhie, J. L. Rocha, F. Ryabov, O. A. Ryder, S. Sacco, K. Shafin, V. A. Shepelev, V. Slon, S. J. Solar, J. M. Storer, P. H. Sudmant, Sweetalana, A. Sweeten, M. G. Tassia, F. Thibaud-Nissen, M. Ventura, M. A. Wilson, A. C. Young, H. Zeng, X. Zhang, Z. A. Szpiech, C. D. Huber, J. L. Gerton, S. V. Yi, M. C. Schatz, I. A. Alexandrov, S. Koren, R. J. O’Neill, E. E. Eichler, A. M. Phillippy, The complete sequence and comparative analysis of ape sex chromosomes. Nature 630, 401–411 (2024).

16. D. Yoo, A. Rhie, P. Hebbar, F. Antonacci, G. A. Logsdon, S. J. Solar, D. Antipov, B. D. Pickett, Y. Safonova, F. Montinaro, Y. Luo, J. Malukiewicz, J. M. Storer, J. Lin, A. N. Sequeira, R. J. Mangan, G. Hickey, G. M. Anez, P. Balachandran, A. Bankevich, C. R. Beck, A. Biddanda, M. Borchers, G. G. Bouffard, E. Brannan, S. Y. Brooks, L. Carbone, L. Carrel, A. P. Chan, J. Crawford, M. Diekhans, E. Engelbrecht, C. Feschotte, G. Formenti, G. H. Garcia, L. de Gennaro, D. Gilbert, R. E. Green, A. Guarracino, I. Gupta, D. Haddad, J. Han, R. S. Harris, G. A. Hartley, W. T. Harvey, M. Hiller, K. Hoekzema, M. L. Houck, H. Jeong, K. Kamali, M. Kellis, B. Kille, C. Lee, Y. Lee, W. Lees, A. P. Lewis, Q. Li, M. Loftus, Y. H. E. Loh, H. Loucks, J. Ma, Y. Mao, J. F. I. Martinez, P. Masterson, R. C. McCoy, B. McGrath, S. McKinney, B. S. Meyer, K. H. Miga, S. K. Mohanty, K. M. Munson, K. Pal, M. Pennell, P. A. Pevzner, D. Porubsky, T. Potapova, F. R. Ringeling, J. L. Rocha, O. A. Ryder, S. Sacco, S. Saha, T. Sasaki, M. C. Schatz, N. J. Schork, C. Shanks, L. Smeds, D. R. Son, C. Steiner, A. P. Sweeten, M. G. Tassia, F. Thibaud-Nissen, E. Torres-González, M. Trivedi, W. Wei, J. Wertz, M. Yang, P. Zhang, S. Zhang, Y. Zhang, Z. Zhang, S. A. Zhao, Y. Zhu, E. D. Jarvis, J. L. Gerton, I. Rivas-González, B. Paten, Z. A. Szpiech, C. D. Huber, T. L. Lenz, M. K. Konkel, S. V. Yi, S. Canzar, C. T. Watson, P. H. Sudmant, E. Molloy, E. Garrison, C. B. Lowe, M. Ventura, R. J. O’Neill, S. Koren, K. D. Makova, A. M. Phillippy, E. E. Eichler, Complete sequencing of ape genomes. Nature 641, 401–418 (2025).

17. H. S. Malik, S. Henikoff, Major evolutionary transitions in centromere complexity. Cell 138, 1067–1082 (2009).

18. K. H. Miga, I. A. Alexandrov, Variation and evolution of human centromeres: A field guide and perspective. Annu. Rev. Genet. 55, 583–602 (2021).

19. S. Henikoff, K. Ahmad, H. S. Malik, The centromere paradox: stable inheritance with rapidly evolving DNA. Science 293, 1098–1102 (2001).

20. C. Courret, L. W. Hemmer, X. Wei, P. D. Patel, B. J. Chabot, N. J. Fuda, X. Geng, C.-H. Chang, B. G. Mellone, A. M. Larracuente, Turnover of retroelements and satellite DNA drives centromere reorganization over short evolutionary timescales in Drosophila. PLoS Biol. 22, e3002911 (2024).

21. T. Fukagawa, W. C. Earnshaw, The centromere: chromatin foundation for the kinetochore machinery. Dev. Cell 30, 496–508 (2014).

22. K. L. McKinley, I. M. Cheeseman, The molecular basis for centromere identity and function. Nat. Rev. Mol. Cell Biol. 17, 16–29 (2016).

23. B. E. Black, D. W. Cleveland, Epigenetic centromere propagation and the nature of CENP-a nucleosomes. Cell 144, 471–479 (2011).

24. R. C. Allshire, G. H. Karpen, Epigenetic regulation of centromeric chromatin: old dogs, new tricks? Nat. Rev. Genet. 9, 923–937 (2008).

25. C. Salinas-Luypaert, D. Dubocanin, R. J. Lee, L. Andrade Ruiz, R. Gamba, M. Grison, L. Velikovsky, A. Angrisani, A. Scelfo, Y. Xu, M. Dumont, V. Barra, T. Wilhelm, G. Velasco, M. Losito, R. Wardenaar, C. Francastel, F. Foijer, G. J. P. L. Kops, K. H. Miga, N. Altemose, D. Fachinetti, DNA methylation influences human centromere positioning and function. Nat. Genet. 57, 2509–2521 (2025).

26. P. Guckelberger, L. Haut, R. Tornisiello, H. Kretzmer, A. Meissner, HELLS is required for maintaining proper DNA modification at human satellite repeats. Genome Biol. 26, 211 (2025).

27. J. D. Lewis, R. R. Meehan, W. J. Henzel, I. Maurer-Fogy, P. Jeppesen, F. Klein, A. Bird, Purification, sequence, and cellular localization of a novel chromosomal protein that binds to methylated DNA. Cell 69, 905–914 (1992).

28. I. Mendizabal, S. V. Yi, Whole-genome bisulfite sequencing maps from multiple human tissues reveal novel CpG islands associated with tissue-specific regulation. Hum. Mol. Genet. 25, 69–82 (2016).

29. Consortium, The ENCODE Project, An integrated encyclopedia of DNA elements in the human genome. Nature 489, 57–74 (2012).

30. I. Mendizabal, S. Berto, N. Usui, K. Toriumi, P. Chatterjee, C. Douglas, I. Huh, H. Jeong, T. Layman, C. A. Tamminga, T. M. Preuss, G. Konopka, S. V. Yi, Cell type-specific epigenetic links to schizophrenia risk in the brain. Genome Biol. 20, 135 (2019).

31. H. E. Zimmerman, J. L. Moore, R. H. Miller, I. Stirland, A. Jenkins, E. Saito, T. Jenkins, J. T. Hill, ModSeqR: an R package for efficient analysis of modified nucleotide data. BMC Genomics 27 (2026).

32. N. Elango, S. V. Yi, DNA methylation and structural and functional bimodality of vertebrate promoters. Mol. Biol. Evol. 25, 1602–1608 (2008).

33. M. Weber, J. J. Davies, D. Wittig, E. J. Oakeley, M. Haase, W. L. Lam, D. Schubeler, Chromosome-wide and promoter-specific analyses identify sites of differential DNA methylation in normal and transformed human cells. Nat. Genet. 37, 853–862 (2005).

34. B. Yu, X. Dong, S. Gravina, Ö. Kartal, T. Schimmel, J. Cohen, D. Tortoriello, R. Zody, R. D. Hawkins, J. Vijg, Genome-wide, single-cell DNA methylomics reveals increased non-CpG methylation during human oocyte maturation. Stem Cell Reports 9, 397–407 (2017).

35. D. Yoo, K. M. Munson, E. E. Eichler, Epigenetic and evolutionary features of ape subterminal heterochromatin. Genome Res. 36, 38–49 (2026).

36. D. C. Soto, J. M. Uribe-Salazar, G. Kaya, R. Valdarrago, A. Sekar, N. K. Haghani, K. Hino, G. La, N. A. F. Mariano, C. Ingamells, A. Baraban, Z. Jamal, T. N. Turner, E. D. Green, S. Simó, G. Quon, A. M. Andrés, M. Y. Dennis, Human-specific gene expansions contribute to brain evolution. Cell 188, 5363–5383.e22 (2025).

37. GTEx Consortium, Genetic effects on gene expression across human tissues. Nature 550, 204 (2017).

38. D. A. Glinos, G. Garborcauskas, P. Hoffman, N. Ehsan, L. Jiang, A. Gokden, X. Dai, F. Aguet, K. L. Brown, K. Garimella, T. Bowers, M. Costello, K. Ardlie, R. Jian, N. R. Tucker, P. T. Ellinor, E. D. Harrington, H. Tang, M. Snyder, S. Juul, P. Mohammadi, D. G. MacArthur, T. Lappalainen, B. B. Cummings, Transcriptome variation in human tissues revealed by long-read sequencing. Nature 608, 353–359 (2022).

39. N. Elango, S.-H. Kim, E. Vigoda, S. V. Yi, Mutations of different molecular origins exhibit contrasting patterns of regional substitution rate variation. PLoS Comput. Biol. 4, e1000015 (2008).

40. S.-H. Kim, N. Elango, C. Warden, E. Vigoda, S. V. Yi, Heterogeneous genomic molecular clocks in primates. PLoS Genet. 2, e163 (2006).

41. K. Yamagata, T. Yamazaki, H. Miki, N. Ogonuki, K. Inoue, A. Ogura, T. Baba, Centromeric DNA hypomethylation as an epigenetic signature discriminates between germ and somatic cell lineages. Dev. Biol. 312, 419–426 (2007).

42. D. Porubsky, H. Dashnow, T. A. Sasani, G. A. Logsdon, P. Hallast, M. D. Noyes, Z. N. Kronenberg, T. Mokveld, N. Koundinya, C. Nolan, C. J. Steely, A. Guarracino, E. Dolzhenko, W. T. Harvey, W. J. Rowell, K. Grigorev, T. J. Nicholas, M. E. Goldberg, K. K. Oshima, J. Lin, P. Ebert, W. S. Watkins, T. Y. Leung, V. C. T. Hanlon, S. McGee, B. S. Pedersen, H. C. Happ, H. Jeong, K. M. Munson, K. Hoekzema, D. D. Chan, Y. Wang, J. Knuth, G. H. Garcia, C. Fanslow, C. Lambert, C. Lee, J. D. Smith, S. Levy, C. E. Mason, E. Garrison, P. M. Lansdorp, D. W. Neklason, L. B. Jorde, A. R. Quinlan, M. A. Eberle, E. E. Eichler, Human de novo mutation rates from a four-generation pedigree reference. Nature 643, 427–436 (2025).

43. J. Lin, F. K. Mastrorosa, M. D. Noyes, D. Yoo, A. Rhie, D. Porubsky, K. Hoekzema, K. M. Munson, N. Koundinya, W. S. Watkins, L. B. Jorde, A. R. Quinlan, D. W. Neklason, A. M. Phillippy, E. E. Eichler, Human acrocentric chromosome short-arm de novo mutation and recombination. Cell 189, 4876–4890.e8 (2026).

44. X. She, J. E. Horvath, Z. Jiang, G. Liu, T. S. Furey, L. Christ, R. Clark, T. Graves, C. L. Gulden, C. Alkan, J. A. Bailey, C. Sahinalp, M. Rocchi, D. Haussler, R. K. Wilson, W. Miller, S. Schwartz, E. E. Eichler, The structure and evolution of centromeric transition regions within the human genome. Nature 430, 857–864 (2004).

45. H. C. Mefford, B. J. Trask, The complex structure and dynamic evolution of human subtelomeres. Nat. Rev. Genet. 3, 91–102 (2002).

46. E. V. Linardopoulou, E. M. Williams, Y. Fan, C. Friedman, J. M. Young, B. J. Trask, Human subtelomeres are hot spots of interchromosomal recombination and segmental duplication. Nature 437, 94–100 (2005).

47. H. Kaessmann, Origins, evolution, and phenotypic impact of new genes. Genome Res. 20, 1313–1326 (2010).

48. E. Witt, S. Benjamin, N. Svetec, L. Zhao, Testis single-cell RNA-seq reveals the dynamics of de novo gene transcription and germline mutational bias in Drosophila. Elife 8, e47138 (2019).

49. M. Soumillon, A. Necsulea, M. Weier, D. Brawand, X. Zhang, H. Gu, P. Barthès, M. Kokkinaki, S. Nef, A. Gnirke, M. Dym, B. de Massy, T. S. Mikkelsen, H. Kaessmann, Cellular source and mechanisms of high transcriptome complexity in the mammalian testis. Cell Rep. 3, 2179–2190 (2013).

50. B. Xia, Y. Yan, M. Baron, F. Wagner, D. Barkley, M. Chiodin, S. Y. Kim, D. L. Keefe, J. P. Alukal, J. D. Boeke, I. Yanai, Widespread transcriptional scanning in the testis modulates gene evolution rates. Cell 180, 248–262.e21 (2020).

51. S. K. Reilly, J. Yin, A. E. Ayoub, D. Emera, J. Leng, J. Cotney, R. Sarro, P. Rakic, J. P. Noonan, Evolutionary genomics. Evolutionary changes in promoter and enhancer activity during human corticogenesis. *Science (New York*, N.Y.) 347, 1155–1159 (2015).

52. M. W. Vermunt, S. C. Tan, B. Castelijns, G. Geeven, P. Reinink, E. de Bruijn, I. Kondova, S. Persengiev, R. Bontrop, E. Cuppen, W. de Laat, M. P. Creyghton, B. Netherlands Brain, Epigenomic annotation of gene regulatory alterations during evolution of the primate brain. Nature Neuroscience 19, 494–503 (2016).

53. H. Won, J. Huang, C. K. Opland, C. L. Hartl, D. H. Geschwind, Human evolved regulatory elements modulate genes involved in cortical expansion and neurodevelopmental disease susceptibility. Nat. Commun. 10, 2396 (2019).

54. J. H. Notwell, T. Chung, W. Heavner, G. Bejerano, A family of transposable elements co-opted into developmental enhancers in the mouse neocortex. Nat. Commun. 6, 6644 (2015).

55. A. Y. Du, J. D. Chobirko, X. Zhuo, C. Feschotte, T. Wang, Regulatory transposable elements in the encyclopedia of DNA elements. Nat. Commun. 15, 7594 (2024).

56. M. Trizzino, Y. Park, M. Holsbach-Beltrame, K. Aracena, K. Mika, M. Caliskan, G. H. Perry, V. J. Lynch, C. D. Brown, Transposable elements are the primary source of novelty in primate gene regulation. Genome Res. 27, 1623–1633 (2017).

57. H. Sasaki, Y. Matsui, Epigenetic events in mammalian germ-cell development: reprogramming and beyond. Nat. Rev. Genet. 9, 129–140 (2008).

58. W. Reik, W. Dean, J. Walter, Epigenetic Reprogramming in Mammalian Development. Science 293, 1089–1093 (2001).

59. D. A. Gallegos, U. Chan, L.-F. Chen, A. E. West, Chromatin regulation of neuronal maturation and plasticity. Trends Neurosci. 41, 311–324 (2018).

60. Y. Su, J. Shin, C. Zhong, S. Wang, P. Roychowdhury, J. Lim, D. Kim, G.-L. Ming, H. Song, Neuronal activity modifies the chromatin accessibility landscape in the adult brain. Nat. Neurosci. 20, 476–483 (2017).

61. L. de la Torre-Ubieta, J. L. Stein, H. Won, C. K. Opland, D. Liang, D. Lu, D. H. Geschwind, The dynamic landscape of open chromatin during human cortical neurogenesis. Cell 172, 289–304.e18 (2018).

62. S. Seisenberger, S. Andrews, F. Krueger, J. Arand, J. Walter, F. Santos, C. Popp, B. Thienpont, W. Dean, W. Reik, The dynamics of genome-wide DNA methylation reprogramming in mouse primordial germ cells. Mol. Cell 48, 849–862 (2012).

63. R. Lister, E. A. Mukamel, J. R. Nery, M. Urich, C. A. Puddifoot, N. D. Johnson, J. Lucero, Y. Huang, A. J. Dwork, M. D. Schultz, M. Yu, J. Tonti-Filippini, H. Heyn, S. Hu, J. C. Wu, A. Rao, M. Esteller, C. He, F. G. Haghighi, T. J. Sejnowski, M. M. Behrens, J. R. Ecker, Global Epigenomic Reconfiguration During Mammalian Brain Development. Science 341, 1237905 (2013).

64. C. Roadmap Epigenomics, A. Kundaje, W. Meuleman, J. Ernst, M. Bilenky, A. Yen, A. Heravi-Moussavi, P. Kheradpour, Z. Zhang, J. Wang, M. J. Ziller, V. Amin, J. W. Whitaker, M. D. Schultz, L. D. Ward, A. Sarkar, G. Quon, R. S. Sandstrom, M. L. Eaton, Y.-C. Wu, A. R. Pfenning, X. Wang, M. Claussnitzer, Y. Liu, C. Coarfa, R. A. Harris, N. Shoresh, C. B. Epstein, E. Gjoneska, D. Leung, W. Xie, R. D. Hawkins, R. Lister, C. Hong, P. Gascard, A. J. Mungall, R. Moore, E. Chuah, A. Tam, T. K. Canfield, R. S. Hansen, R. Kaul, P. J. Sabo, M. S. Bansal, A. Carles, J. R. Dixon, K.-H. Farh, S. Feizi, R. Karlic, A.-R. Kim, A. Kulkarni, D. Li, R. Lowdon, G. Elliott, T. R. Mercer, S. J. Neph, V. Onuchic, P. Polak, N. Rajagopal, P. Ray, R. C. Sallari, K. T. Siebenthall, N. A. Sinnott-Armstrong, M. Stevens, R. E. Thurman, J. Wu, B. Zhang, X. Zhou, A. E. Beaudet, L. A. Boyer, P. L. De Jager, P. J. Farnham, S. J. Fisher, D. Haussler, S. J. M. Jones, W. Li, M. A. Marra, M. T. McManus, S. Sunyaev, J. A. Thomson, T. D. Tlsty, L.-H. Tsai, W. Wang, R. A. Waterland, M. Q. Zhang, L. H. Chadwick, B. E. Bernstein, J. F. Costello, J. R. Ecker, M. Hirst, A. Meissner, A. Milosavljevic, B. Ren, J. A. Stamatoyannopoulos, T. Wang, M. Kellis, Integrative analysis of 111 reference human epigenomes. Nature 518, 317–330 (2015).

65. M. Kim, J. Costello, DNA methylation: an epigenetic mark of cellular memory. Exp. Mol. Med. 49, e322 (2017).

66. K. D. Hansen, S. Sabunciyan, B. Langmead, N. Nagy, R. Curley, G. Klein, E. Klein, D. Salamon, A. P. Feinberg, Large-scale hypomethylated blocks associated with Epstein-Barr virus-induced B-cell immortalization. Genome Res. 24, 177–184 (2014).

67. X. Chen, Q. Lin, J. Wen, W. Lin, J. Liang, H. Huang, L. Li, J. Huang, F. Chen, D. Liu, G. Chen, Whole genome bisulfite sequencing of human spermatozoa reveals differentially methylated patterns from type 2 diabetic patients. J. Diabetes Investig. 11, 856–864 (2020).

68. F. Krueger, F. James, P. Ewels, E. Afyounian, M. Weinstein, B. Schuster-Boeckler, G. Hulselmans, sclamons, *FelixKrueger/TrimGalore: v0.6.10 - Add Default Decompression Path* (Zenodo, 2023; 10.5281/zenodo.7598955).

69. F. Krueger, S. R. Andrews, Bismark: a flexible aligner and methylation caller for Bisulfite-Seq applications. Bioinformatics 27, 1571–1572 (2011).

70. A. M. Mc Cartney, K. Shafin, M. Alonge, A. V. Bzikadze, G. Formenti, A. Fungtammasan, K. Howe, C. Jain, S. Koren, G. A. Logsdon, K. H. Miga, A. Mikheenko, B. Paten, A. Shumate, D. C. Soto, I. Sović, J. M. D. Wood, J. M. Zook, A. M. Phillippy, A. Rhie, Chasing perfection: validation and polishing strategies for telomere-to-telomere genome assemblies. Nat. Methods 19, 687–695 (2022).

71. A. Rhie, S. Nurk, M. Cechova, S. J. Hoyt, D. J. Taylor, N. Altemose, P. W. Hook, S. Koren, M. Rautiainen, I. A. Alexandrov, J. Allen, M. Asri, A. V. Bzikadze, N.-C. Chen, C.-S. Chin, M. Diekhans, P. Flicek, G. Formenti, A. Fungtammasan, C. Garcia Giron, E. Garrison, A. Gershman, J. L. Gerton, P. G. S. Grady, A. Guarracino, L. Haggerty, R. Halabian, N. F. Hansen, R. Harris, G. A. Hartley, W. T. Harvey, M. Haukness, J. Heinz, T. Hourlier, R. M. Hubley, S. E. Hunt, S. Hwang, M. Jain, R. K. Kesharwani, A. P. Lewis, H. Li, G. A. Logsdon, J. K. Lucas, W. Makalowski, C. Markovic, F. J. Martin, A. M. Mc Cartney, R. C. McCoy, J. McDaniel, B. M. McNulty, P. Medvedev, A. Mikheenko, K. M. Munson, T. D. Murphy, H. E. Olsen, N. D. Olson, L. F. Paulin, D. Porubsky, T. Potapova, F. Ryabov, S. L. Salzberg, M. E. G. Sauria, F. J. Sedlazeck, K. Shafin, V. A. Shepelev, A. Shumate, J. M. Storer, L. Surapaneni, A. M. Taravella Oill, F. Thibaud-Nissen, W. Timp, M. Tomaszkiewicz, M. R. Vollger, B. P. Walenz, A. C. Watwood, M. H. Weissensteiner, A. M. Wenger, M. A. Wilson, S. Zarate, Y. Zhu, J. M. Zook, E. E. Eichler, R. J. O’Neill, M. C. Schatz, K. H. Miga, K. D. Makova, A. M. Phillippy, The complete sequence of a human Y chromosome. Nature 621, 344–354 (2023).

72. H. Jeong, P. C. Dishuck, D. Yoo, W. T. Harvey, K. M. Munson, A. P. Lewis, J. Kordosky, G. H. Garcia, Human Genome Structural Variation Consortium (HGSVC), F. Yilmaz, P. Hallast, C. Lee, T. Pastinen, E. E. Eichler, Structural polymorphism and diversity of human segmental duplications. Nat. Genet. 57, 390–401 (2025).

73. H. Li, Minimap2: pairwise alignment for nucleotide sequences. Bioinformatics 34, 3094– 3100 (2018).

74. P. Danecek, J. K. Bonfield, J. Liddle, J. Marshall, V. Ohan, M. O. Pollard, A. Whitwham, T. Keane, S. A. McCarthy, R. M. Davies, H. Li, Twelve years of SAMtools and BCFtools. Gigascience 10 (2021).

75. Y. Liao, G. K. Smyth, W. Shi, featureCounts: an efficient general purpose program for assigning sequence reads to genomic features. Bioinformatics 30, 923–930 (2014).

76. G. Hickey, B. Paten, D. Earl, D. Zerbino, D. Haussler, HAL: a hierarchical format for storing and analyzing multiple genome alignments. Bioinformatics 29, 1341–1342 (2013).

77. L. Laurent, E. Wong, G. Li, T. Huynh, A. Tsirigos, C. T. Ong, H. M. Low, K. W. Kin Sung, I. Rigoutsos, J. Loring, C.-L. Wei, Dynamic changes in the human methylome during differentiation. Genome Res. 20, 320–331 (2010).

78. M. D. Schultz, Y. He, J. W. Whitaker, M. Hariharan, E. A. Mukamel, D. Leung, N. Rajagopal, J. R. Nery, M. A. Urich, H. Chen, S. Lin, Y. Lin, I. Jung, A. D. Schmitt, S. Selvaraj, B. Ren, T. J. Sejnowski, W. Wang, J. R. Ecker, Human body epigenome maps reveal noncanonical DNA methylation variation. Nature 523, 212–216 (2015).

79. M. J. Ziller, H. Gu, F. Muller, J. Donaghey, L. T. Y. Tsai, O. Kohlbacher, P. L. De Jager, E. D. Rosen, D. A. Bennett, B. E. Bernstein, A. Gnirke, A. Meissner, Charting a dynamic DNA methylation landscape of the human genome. Nature 500, 477–481 (2013).

80. I. Hernando-Herraez, H. Heyn, M. Fernandez-Callejo, E. Vidal, H. Fernandez-Bellon, J. Prado-Martinez, A. J. Sharp, M. Esteller, T. Marques-Bonet, The interplay between DNA methylation and sequence divergence in recent human evolution. Nucleic Acids Res. 43, 8204– 8214 (2015).

81. G. Kunde-Ramamoorthy, C. Coarfa, E. Laritsky, N. J. Kessler, R. A. Harris, M. Xu, R. Chen, L. Shen, A. Milosavljevic, R. A. Waterland, Comparison and quantitative verification of mapping algorithms for whole-genome bisulfite sequencing. Nucleic Acids Res. 42, e43 (2014).

82. F. Court, C. Tayama, V. Romanelli, A. Martin-Trujillo, I. Iglesias-Platas, K. Okamura, N. Sugahara, C. Simón, H. Moore, J. V. Harness, H. Keirstead, J. V. Sanchez-Mut, E. Kaneki, P. Lapunzina, H. Soejima, N. Wake, M. Esteller, T. Ogata, K. Hata, K. Nakabayashi, D. Monk, Genome-wide parent-of-origin DNA methylation analysis reveals the intricacies of human imprinting and suggests a germline methylation-independent mechanism of establishment. Genome Res. 24, 554–569 (2014).

83. J. Qu, E. Hodges, A. Molaro, P. Gagneux, M. D. Dean, G. J. Hannon, A. D. Smith, Evolutionary expansion of DNA hypomethylation in the mammalian germline genome. Genome Res. 28, 145–158 (2018).

84. X. Dong, M. R. M. Du, Q. Gouil, L. Tian, J. S. Jabbari, R. Bowden, P. L. Baldoni, Y. Chen, G. K. Smyth, S. L. Amarasinghe, C. W. Law, M. E. Ritchie, Benchmarking long-read RNA-sequencing analysis tools using in silico mixtures. Nat. Methods 20, 1810–1821 (2023).

85. Y. H. Sun, A. Wang, C. Song, G. Shankar, R. K. Srivastava, K. F. Au, X. Z. Li, Single-molecule long-read sequencing reveals a conserved intact long RNA profile in sperm. Nat. Commun. 12, 1361 (2021).

