## Supplementary_Figures for "Germline hypomethylation shapes dynamic CpG reservoirs in ape genomes"

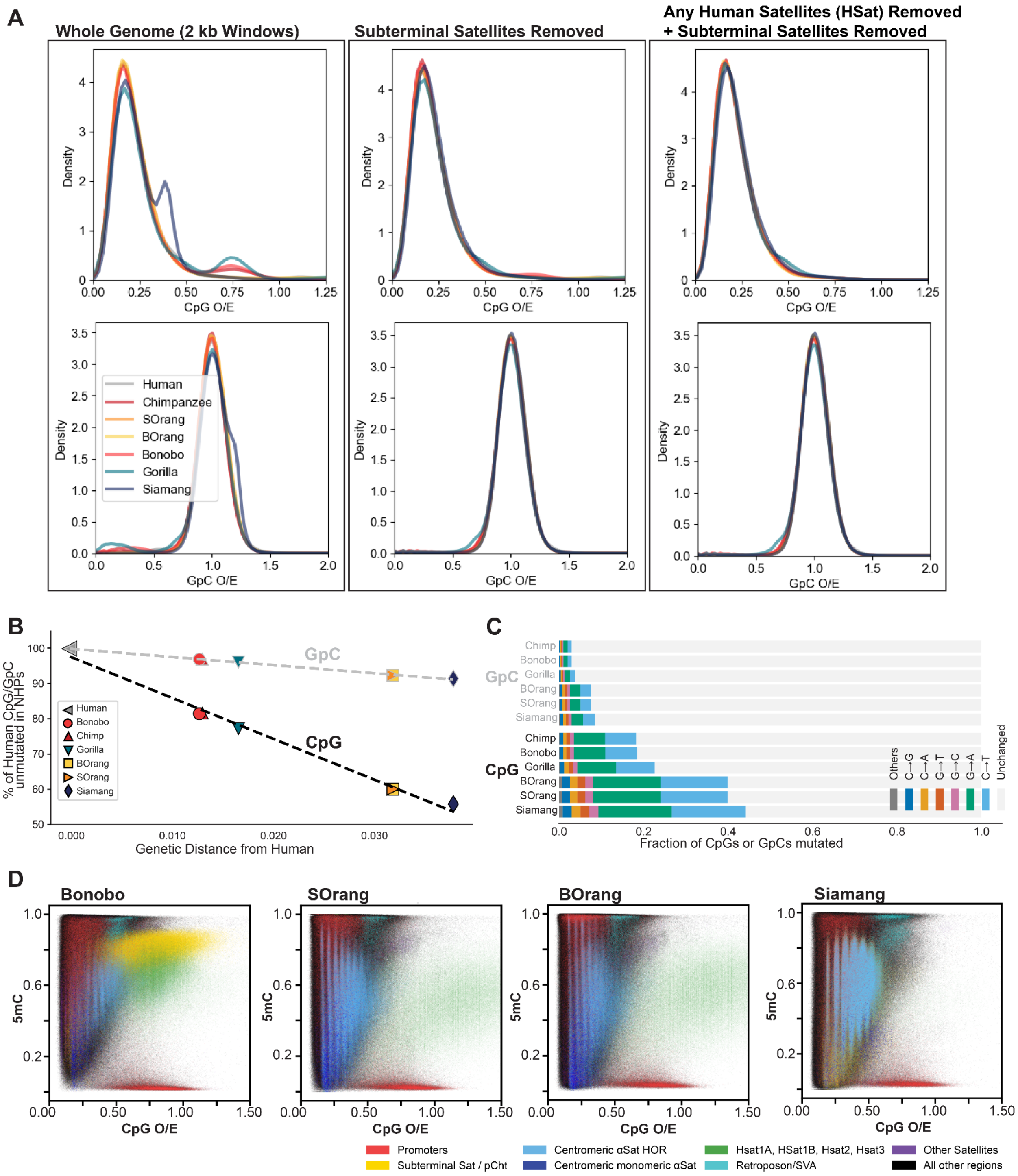


**Fig. S1. Assessment of CpG sites and DNA methylation-driven CpG evolution in ape genomes.** (A) Genome-wide distributions of CpG O/E values in 2-kb windows across complete ape genomes show pervasive CpG depletion. For comparison, GpC O/E values were also calculated in 2-kb windows and are centered around 1, indicating no depletion. Distinct lineage-specific high CpG O/E peaks are observed. These peaks originate from subterminal and HSat satellites; removing windows overlapping subterminal satellites eliminates these peaks, and further removal of HSat windows further removes the remaining peaks. Lineage-expanded

satellite repeats, such as pCht in chimpanzees, gorillas, and bonobos, and subterminal alpha-satellite in siamang form unique CpG-rich peaks. SOrang, Sumatran orangutan; BOrang, Bornean orangutan.
(B-C) Evolutionary dynamics of CpGs and GpCs in human and nonhuman ape genomes. Human CpGs and GpCs were classified as substituted or unchanged at orthologous positions in apes. (B) The number of CpGs conserved across apes decreases rapidly in pairwise comparisons, shown as proportions of total human CpGs. For comparison, the number of conserved GpCs is also shown, revealing a slower decline. (C) Nucleotide substitutions at CpGs and GpCs. CpGs (bottom panel) mutate more rapidly than GpCs, with most mutations at CpGs being transitions associated with deamination of methylated cytosines. Chimp, chimpanzee; SOrang, Sumatran orangutan; BOrang, Bornean orangutan.
(D) T2T ape assemblies were divided into 500-bp windows, and CpG O/E and DNA methylation levels were calculated for each window and plotted to show their relationship. High-coverage somatic DNA methylation profiles were from lymphoblastoid cells for human, chimpanzee, and siamang and from fibroblasts for bonobo, gorilla, Sumatran orangutan, and Bornean orangutan. Each 500-bp window was annotated by genomic feature and color-coded accordingly. Bonobo, orangutans, and siamang are shown here; the other species are shown in Fig. 1D.

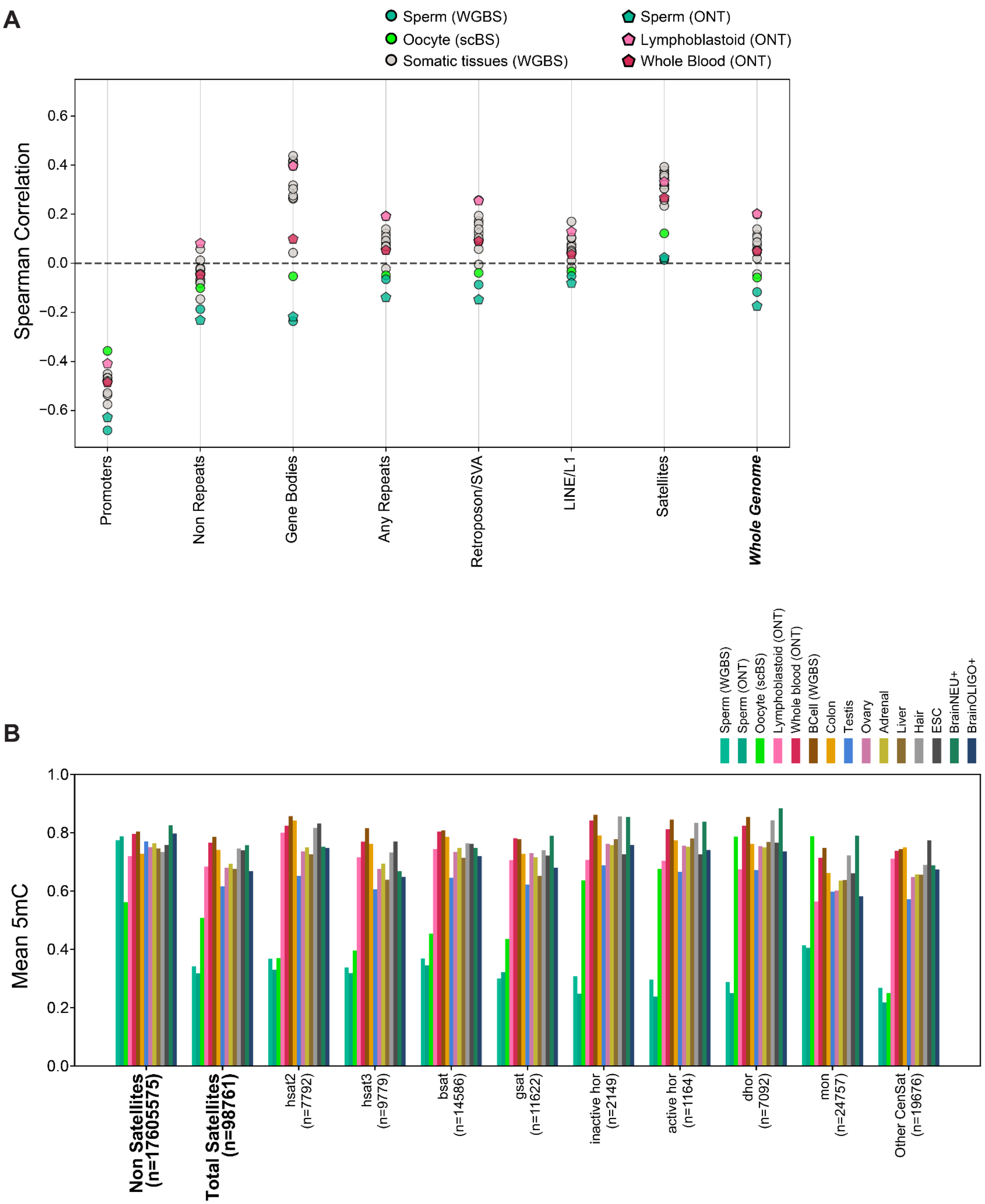


**Fig. S2. Negative correlation between CpG O/E and DNA methylation in oocytes and sperm, and hypomethylation of peri/centromeric satellites.**

(A) Correlations between CpG O/E and DNA methylation were calculated across tissues using shared 500 bp windows for single-cell oocyte methylome samples and other tissues. Oocytes and sperm show generally stronger negative correlations than somatic tissues. The last column shows the genome-wide correlation pattern. (B) Mean DNA methylation levels across satellite and non-satellite regions. Shared CpG sites across tissue datasets, including the aggregated single-cell oocyte dataset, were compared, with the number of CpG sites indicated in parentheses. Oocyte hypomethylation was observed across satellites compared with somatic tissues, with varying degrees among satellite subfamilies. However, these results should be interpreted with caution and further validated using higher-coverage oocyte data. Data are from (*29*, *30*, *34*, *66*, *67*, *77*–*83*), listed in Table S4.

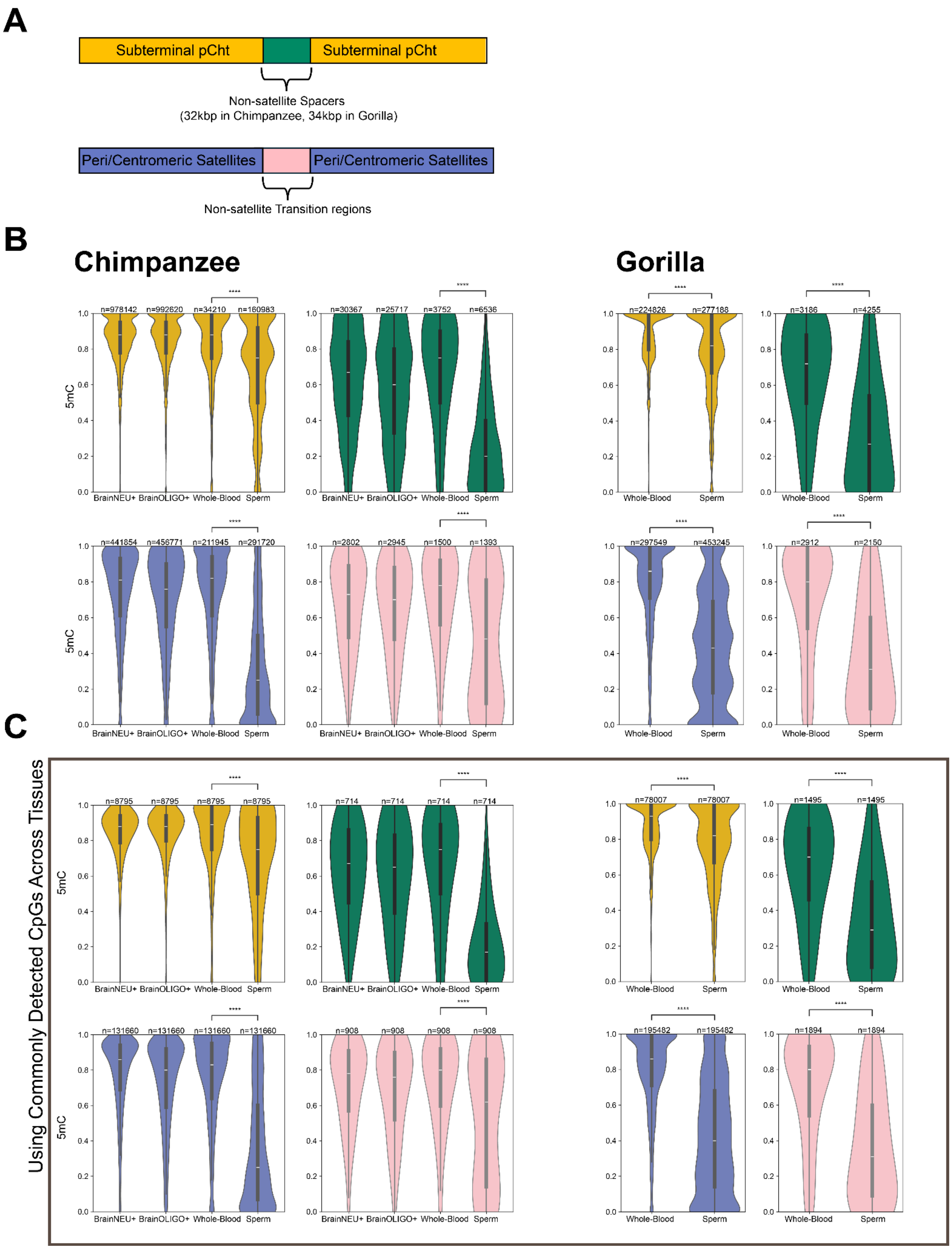


**Fig. S3. DNA methylation levels in subterminal pCht arrays and spacers in somatic tissues and sperm.** (A) Color schematics of the comparison between pCht arrays (yellow) and their spacer regions (green; 32 kb in chimpanzee and 34 kb in gorilla). We also performed an analogous comparison for peri/centromeric satellites (purple) and their embedded non-satellite transition regions (pink), which were sampled to match the size of the pCht spacers (30-34 kb in chimpanzee and 32-36 kb in gorilla). (B) CpG sites were annotated and collected for each region, and their DNA methylation levels were compared across tissues using violin plots. (C) The same analysis was performed using CpG sites commonly detected (shared) across tissues. Two-sided Mann-Whitney-Wilcoxon tests were conducted between whole blood and sperm. ****: $p \leq 1.00e-04$. Pericentromeric satellites are hypomethylated in sperm. Non-satellite regions located between pericentromeric satellites are also hypomethylated. pCht arrays are significantly hypomethylated as well, although not to the same extent as pericentromeric satellites. Spacers embedded within pCht arrays are hypomethylated compared with the pCht arrays themselves, and this pattern is stronger in sperm.

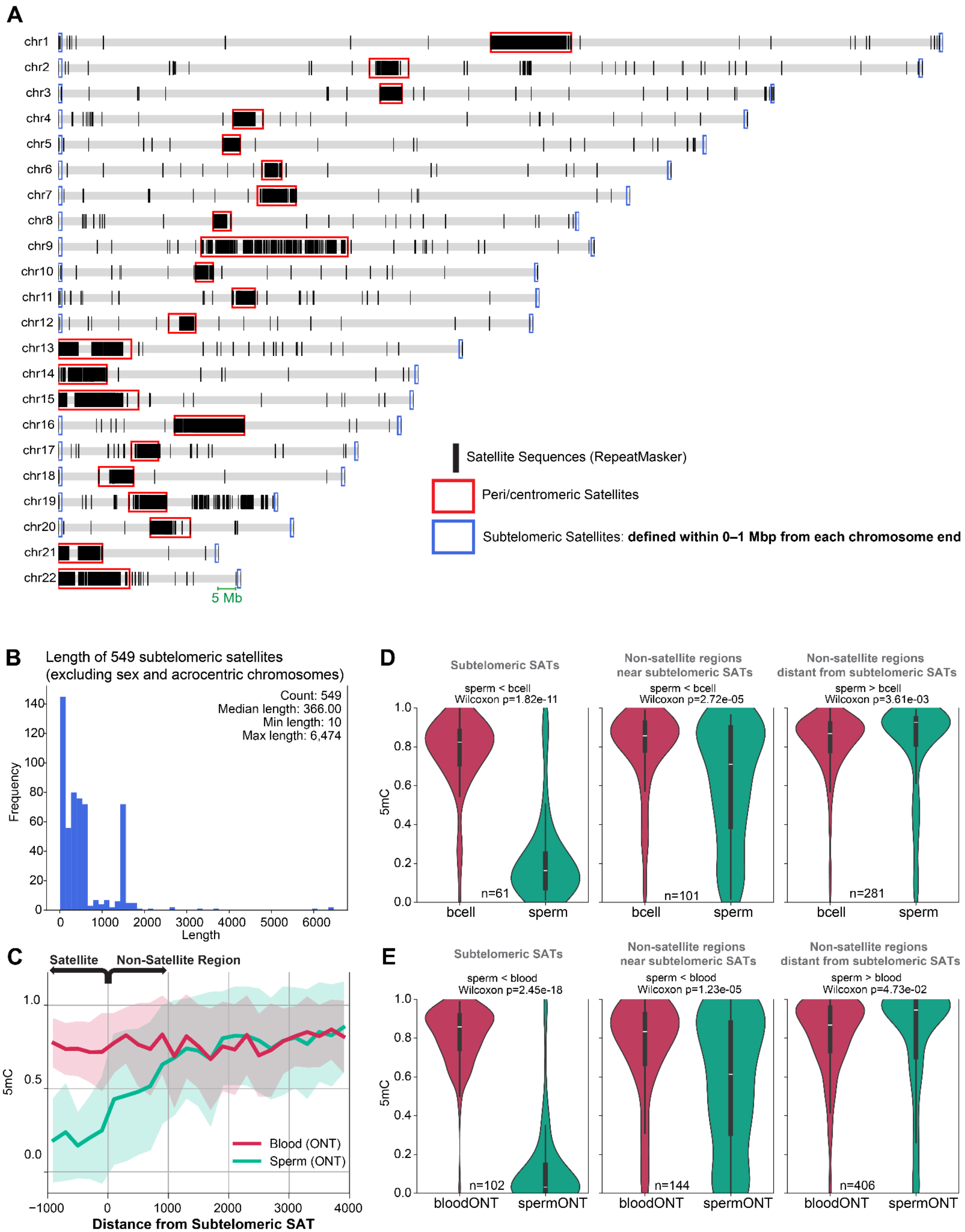


**Fig. S4. Sperm hypomethylation in subtelomeric satellites.** (A) Distribution of satellite sequences, indicated by black bars, in the human genome assembly hs1. Satellite annotations were retrieved from RepeatMasker by selecting annotations that included the term "satellite."

We generated a list of satellites located in subtelomeric regions, defined as regions within 1 Mbp of each chromosome end (blue boxes). Peri/centromeric satellite regions are indicated by red boxes. (B) Length distribution of the 549 identified satellite sequences. Most satellites were short, with a median length of 366 bp and a maximum length of 6,474 bp. (C) DNA methylation landscape near subtelomeric SATs and their adjacent non-satellite regions, comparing ONT-based sperm and blood samples. Negative x-axis values indicate windows within satellite arrays, and positive values indicate non-satellite regions. (D, E) Comparison of DNA methylation levels among subtelomeric SATs, non-satellite regions near subtelomeric SATs (distance: 0–2 kb), and distal non-satellite regions farther from subtelomeric SATs (distance: 5–10 kb). For each category, 500-bp windows were selected based on overlap with or distance from subtelomeric SATs, with the number of windows indicated. DNA methylation levels in B cells and sperm were compared using WGBS data in (D) and ONT data in (E). Subtelomeric SATs were hypomethylated in sperm, and nearby non-satellite regions also showed hypomethylation, indicating that sperm hypomethylation extends into adjacent non-satellite regions. In contrast, distal non-satellite regions did not show sperm hypomethylation. Wilcoxon p-values are indicated.

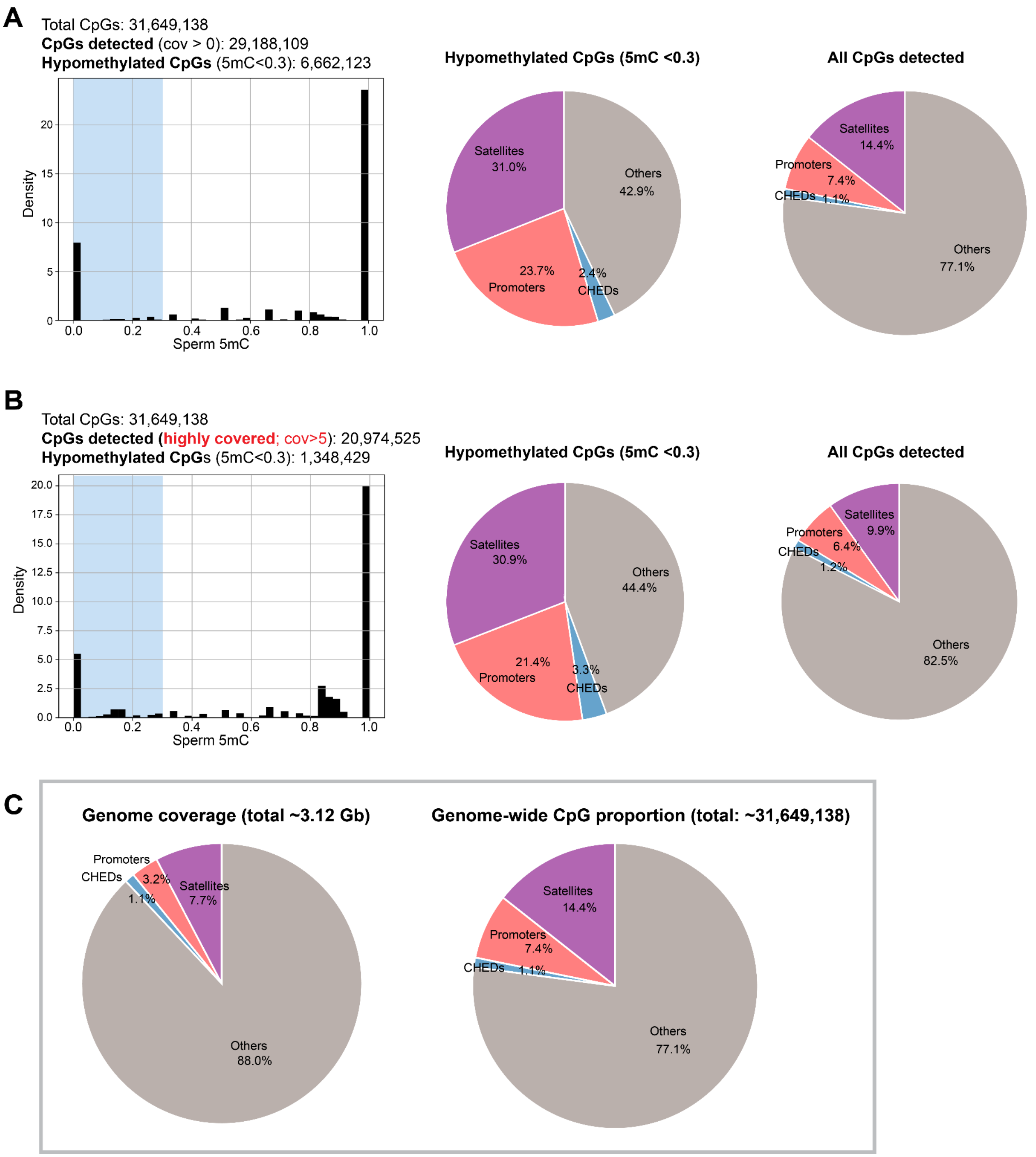


**Fig. S5. Sperm hypomethylated CpG sites are enriched in satellites.** (A) Distribution of CpG DNA methylation levels from ONT sperm data (left). 32.4% of all hypomethylated CpGs (defined as 5mC < 0.3, indicated by blue shading) are located in satellites. In comparison, 23.3% of hypomethylated CpGs are located in promoters. The corresponding proportions for all detected CpGs (right) confirm high enrichment of hypomethylated CpGs in satellites and promoters ($P < 10^{-10}$ by hypergeometric tests for both). (B) The same comparison performed using highly covered CpG sites with coverage > 5, yielded consistent results. (C) Genome-wide base-pair coverage of satellites, promoters, and CHED regions, calculated as the number of base pairs in each category divided by the total genome size. The genome-wide CpG proportion was calculated similarly, as the number of CpGs in each category divided by the total number of genome-wide CpGs. Promoter CpGs within CHED regions are annotated as CHED. Since CHEDs are defined only for autosomes, all comparisons are restricted to autosomal regions.

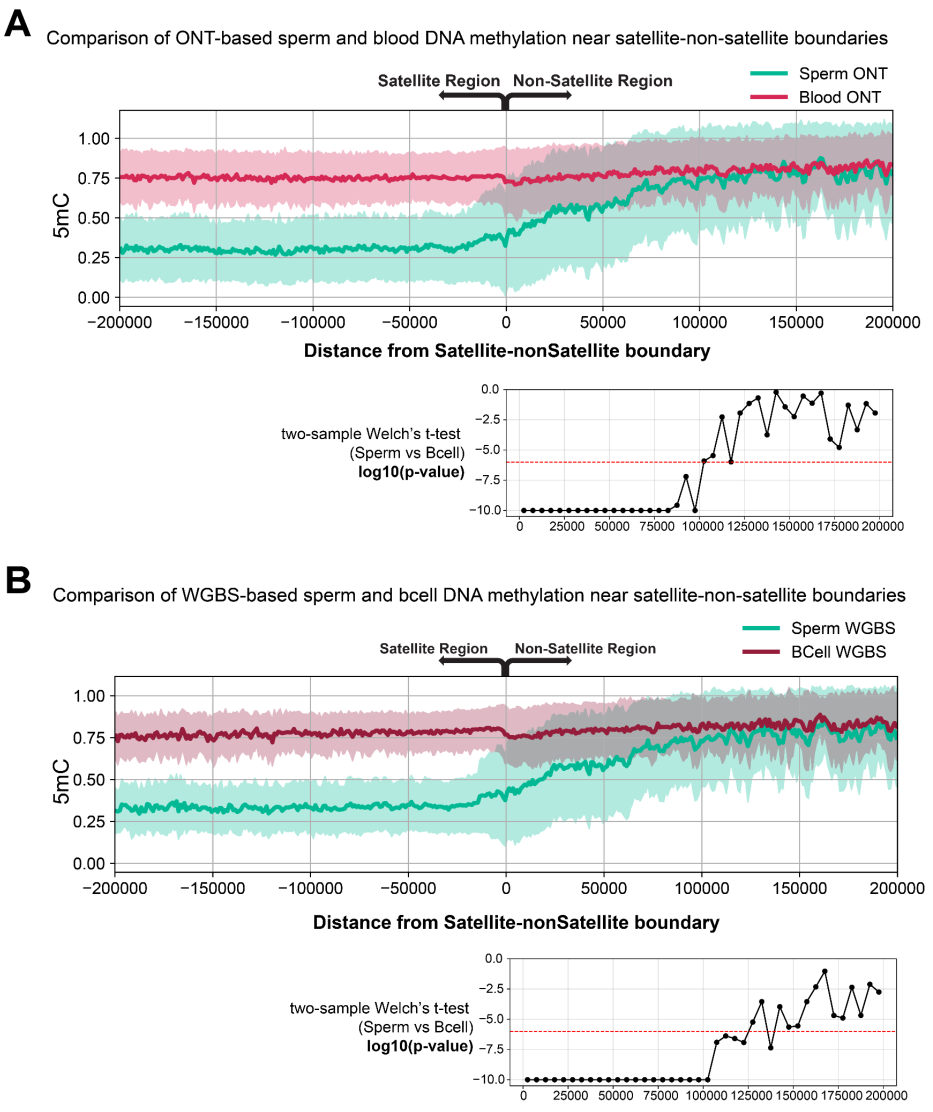


**Fig. S6. Satellite hypomethylation extends into neighboring non-satellite sequences in both ONT and WGBS data.** Sperm shows consistent hypomethylation within satellite arrays, which gradually transitions to methylation levels similar to those of blood/B cells in nearby non-satellite regions. (A) Common 500-bp windows detected in sperm and blood were used to compare DNA methylation landscapes near satellite and non-satellite boundaries. For each window, the distance to the nearest satellite-non-satellite boundary was calculated and associated with DNA methylation patterns. Negative x-axis values indicate windows within satellite arrays, whereas positive x-axis values indicate windows in non-satellite regions, with larger values representing greater distance from the boundary. Sperm hypomethylation remains significant up to ~100 kb, based on a two-sample Welch's t-test using 5-kb sliding windows. (B) The same analysis based on WGBS-based sperm and B cells showed consistent results.

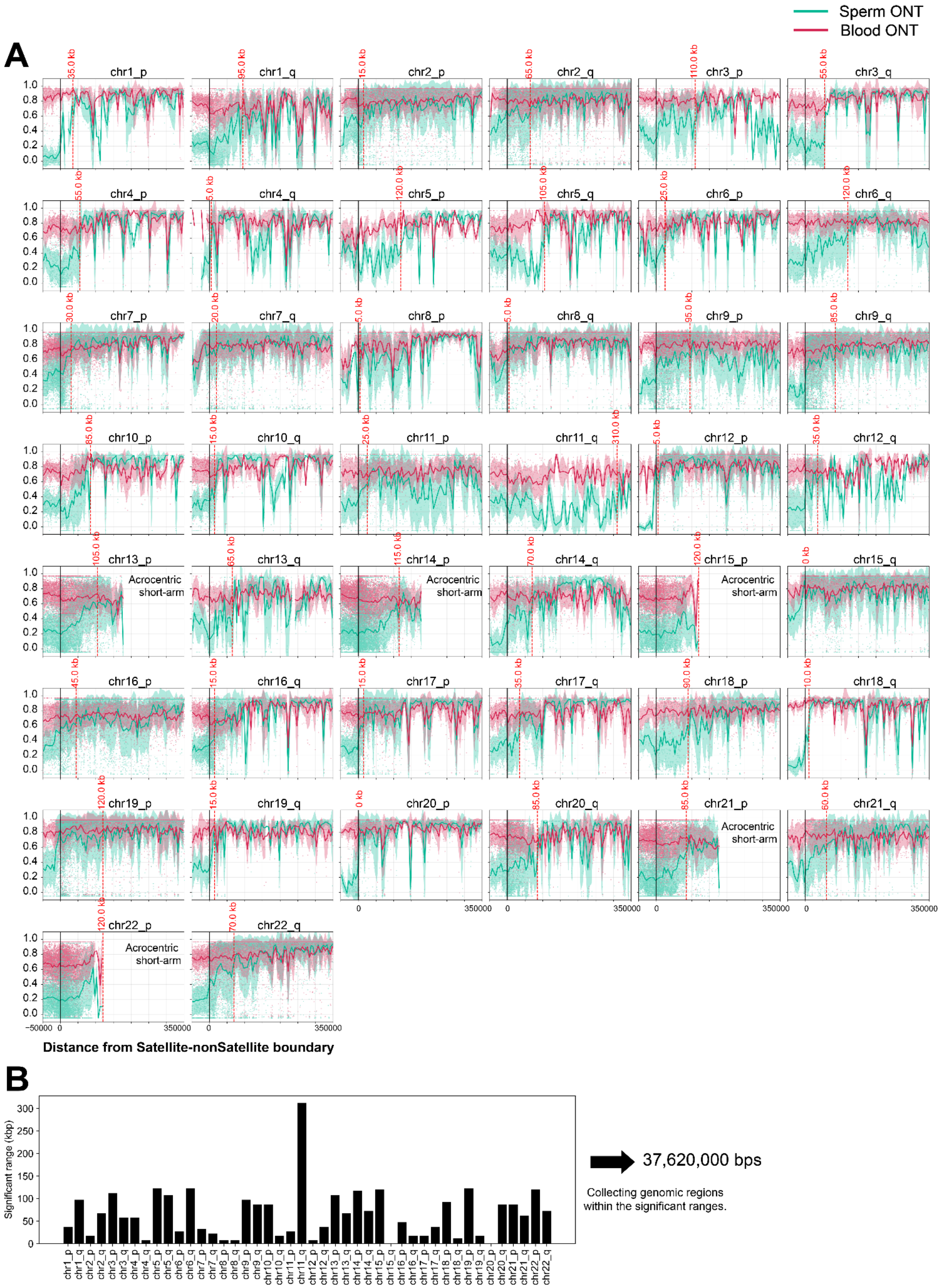


**Fig. S7. Boundaries of centromeric hypomethylation extension domains (CHEDs).** By comparing ONT-based sperm and blood methylation data, we estimated the boundary of significant sperm hypomethylation extending from satellite sequences into nearby non-satellite regions for each chromosome arm. (A) The visualization is analogous to that shown in Fig. S6. We first collected non-satellite windows located within 0 to 10 kb of satellite sequences and tested for sperm hypomethylation using a one-sided Welch's two-sample t-test. We then repeated this test in sliding 10-kb bins with 5-kb overlap, extending up to 400 kb from satellite sequences on each chromosome arm. The range of significant hypomethylation, shown by red lines, was defined as the interval extending from the satellite boundary until two consecutive bins were no longer significant (Benjamini–Hochberg-adjusted FDR $\geq 0.05$). (B) For the defined boundaries, genomic regions within these boundaries were collected for each chromosome, totaling approximately 37.6 Mb.

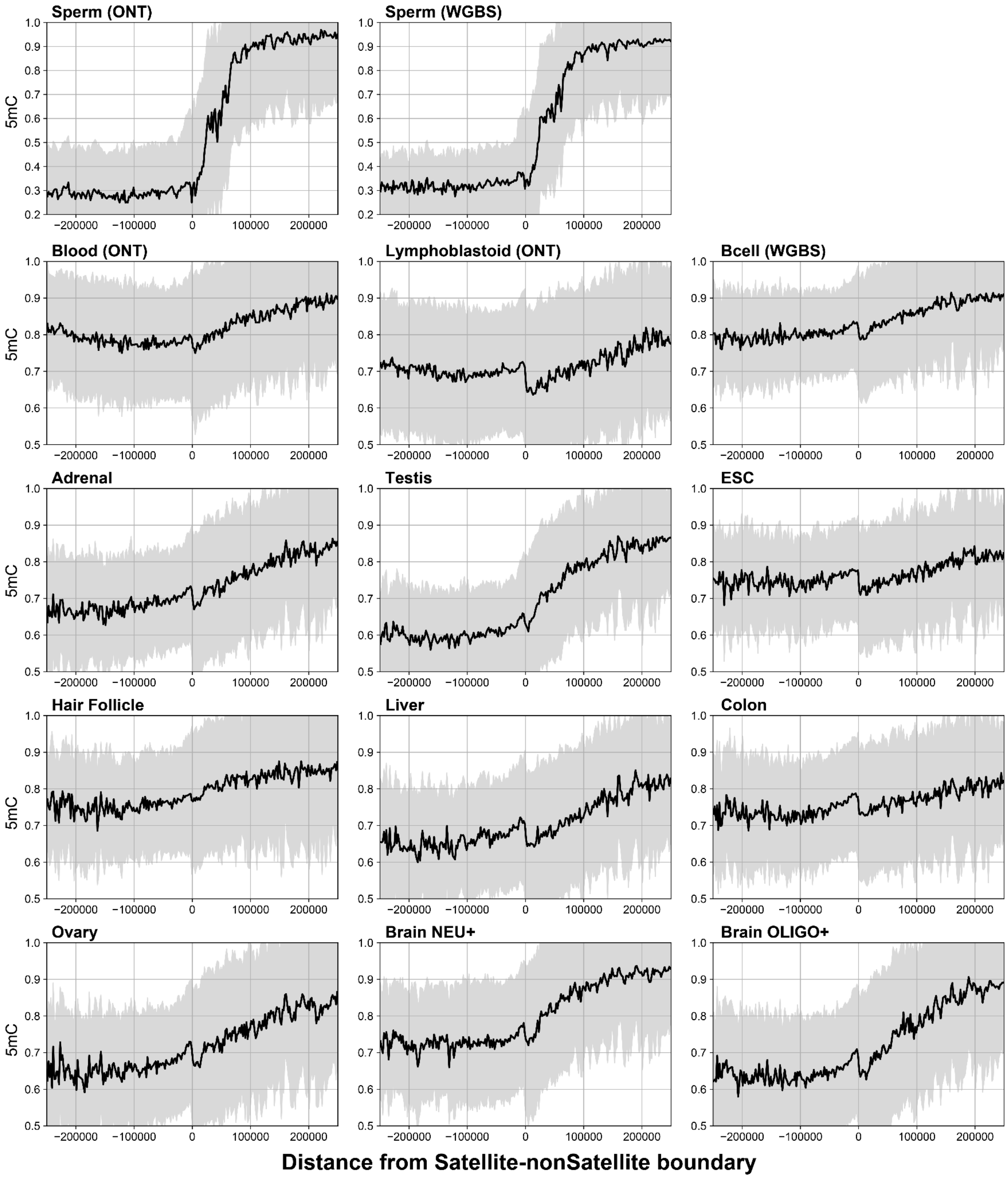


**Fig. S8. Extension of centromeric hypomethylation into adjacent genomic regions across tissues.** DNA methylation landscapes near satellite and non-satellite boundaries across different tissues. The visualization is analogous to that shown in Fig. S6. For each 500-bp window, the distance to the nearest satellite–non-satellite boundary was calculated and associated with DNA methylation patterns. Negative x-axis values indicate windows within satellite arrays, whereas positive x-axis values indicate windows in non-satellite regions, with larger values representing greater distance from the boundary. Tissues including sperm and brain show hypomethylation within satellite arrays to varying degrees, which gradually transitions to higher methylation levels in nearby non-satellite regions, indicating that peri/centromeric methylation patterns extend into adjacent non-satellite regions.

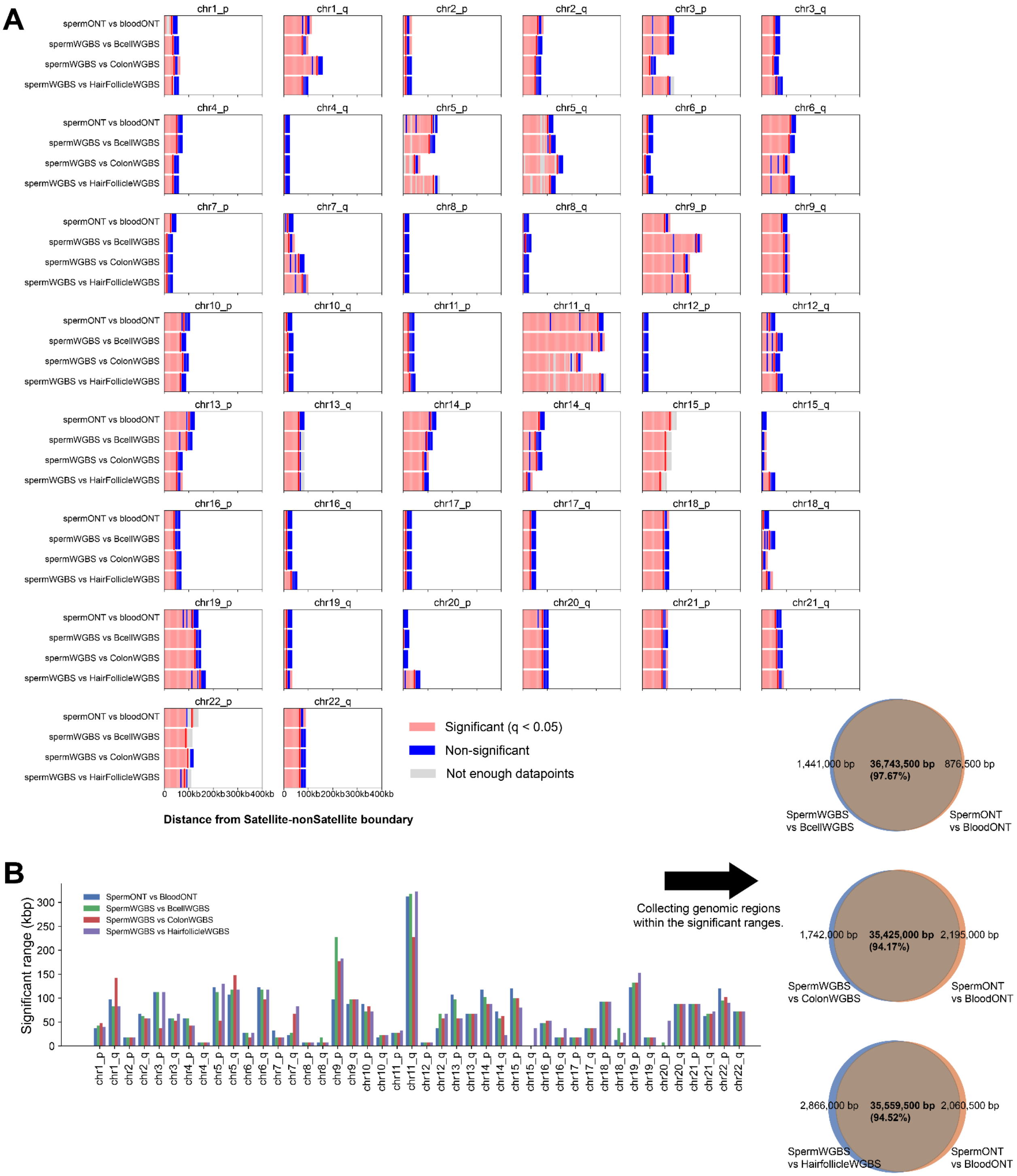


**Fig. S9. Estimation of the boundaries of centromeric hypomethylation extension domains (CHEDs) by comparing sperm methylation profiles with other tissues.** Sperm methylation was compared with tissues lacking extended hypomethylation, including blood, B cells, colon, and hair follicle. (A) The range of significant hypomethylation (red lines) was defined as the interval extending from the satellite boundary until two consecutive bins were no longer significant ($q \geq 0.05$; blue blocks). All pairwise comparisons are shown. We tested for sperm hypomethylation using a one-sided Welch's two-sample t-test (sperm < somatic tissue) in sliding

10-kb bins with 5-kb overlap, extending up to 400 kb from satellite sequences on each chromosome arm. Each block is shown as a rectangle. Sliding bins containing fewer than 10 data points were excluded. P values were corrected for multiple testing using the Benjamini-Hochberg false discovery rate (FDR) procedure.
(B) Genomic regions exhibiting significant hypomethylation were collected, and evaluated between different comparisons. The regions defined in each comparison showed substantial overlap with those identified in the sperm ONT versus blood comparison, with 97.67%, 94.17%, and 94.52% overlap for the sperm versus B cell, colon, and hair follicle comparisons, respectively. A total of 35,033,550 bp was consistently detected across the pairwise comparisons.

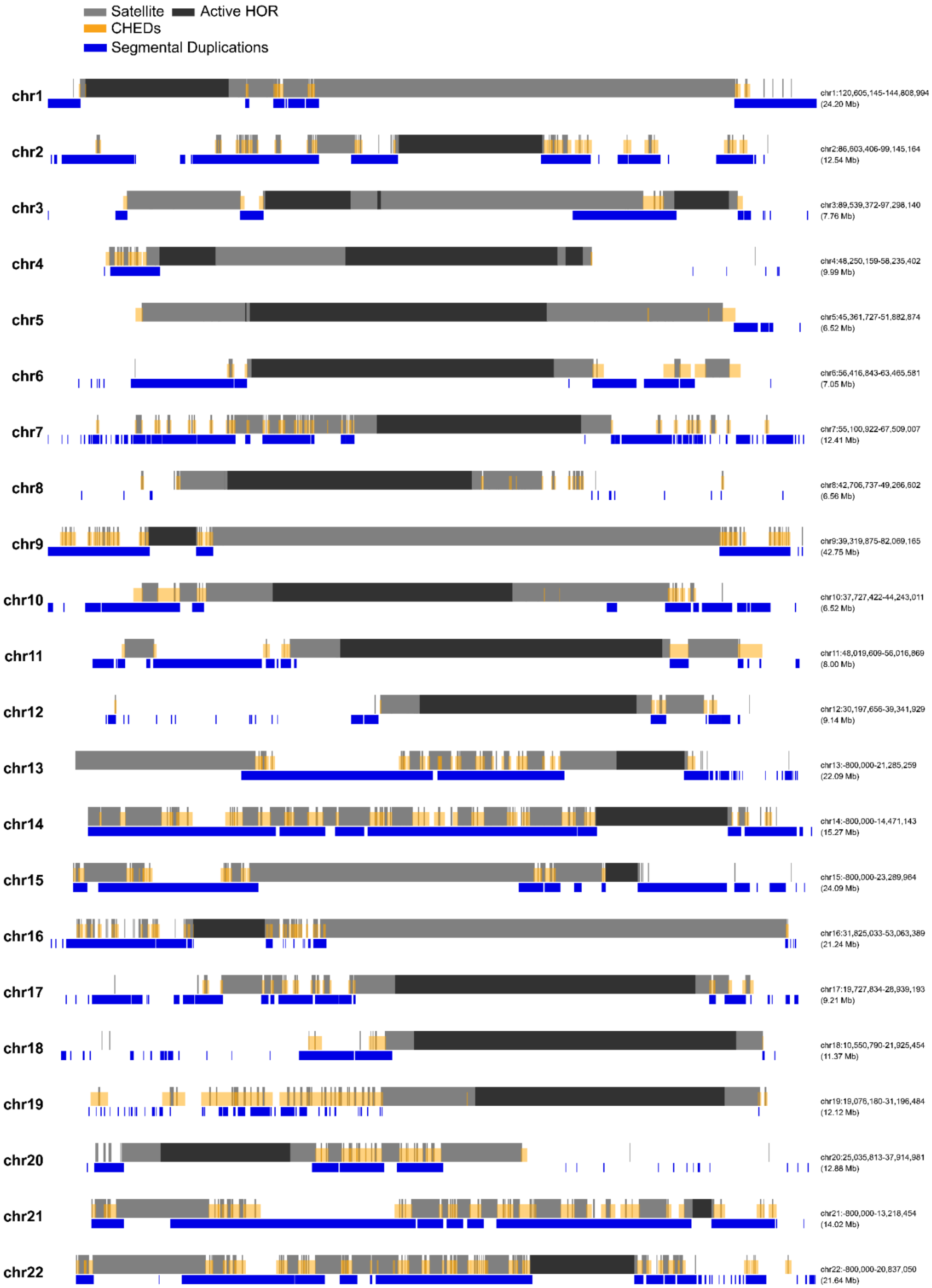


**Fig. S10. Landscape of CHEDs across chromosomes and segmental duplications.** The peri/centromeric region of each chromosome is visualized, with satellite arrays shown in light grey and active centromeric HORs shown in dark grey. CHEDs are indicated by orange boxes. Segmental duplication-associated regions are also indicated by blue boxes below.

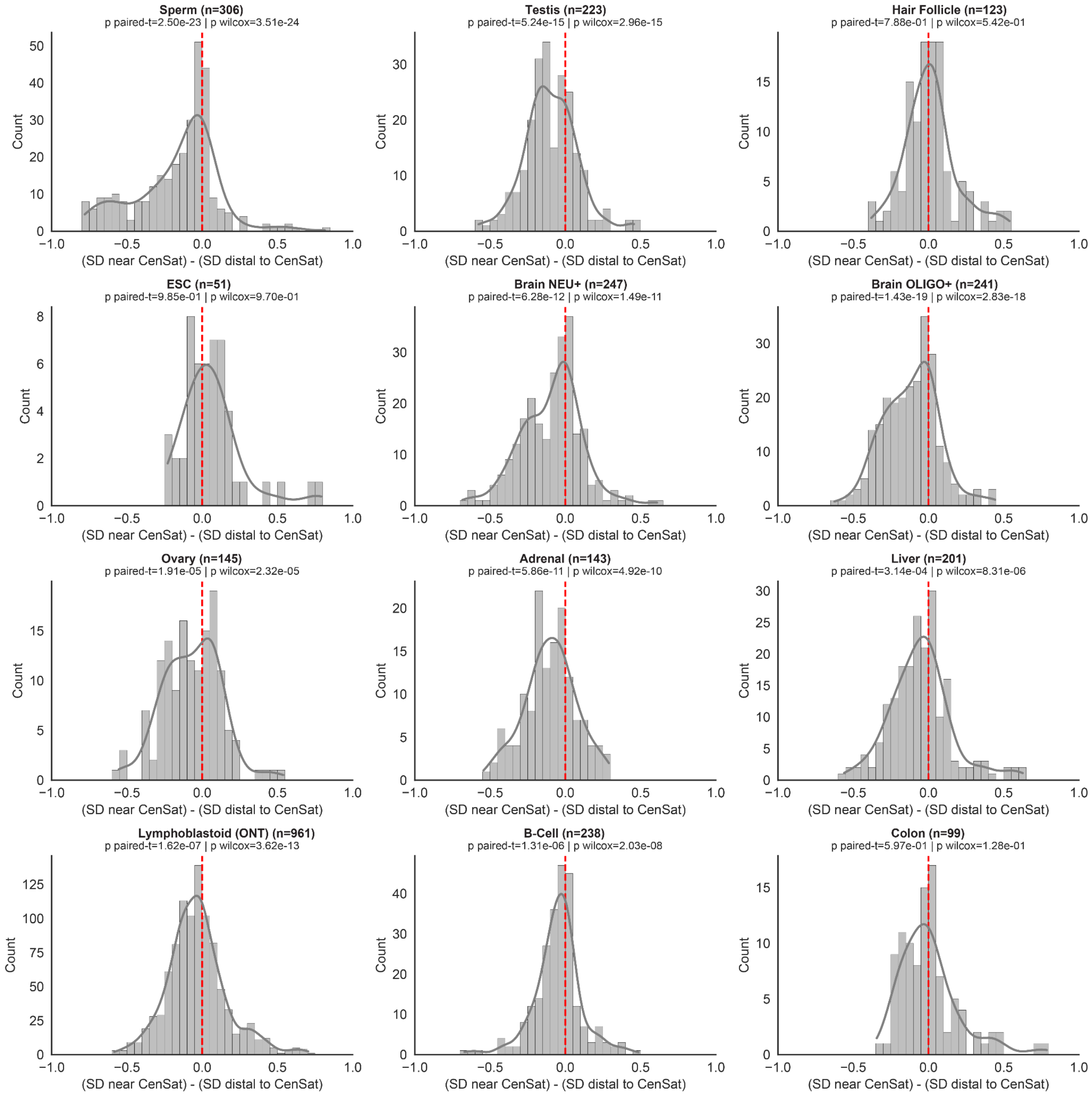


**Fig. S11. Hypomethylation of segmental duplications near pericentromeric regions.** Highly identical segmental duplication (SD; identity >95%) pairs were compared for DNA methylation in each tissue, with one copy located near satellite arrays (SD near CenSat; 0 < distance from satellite array < 100 kb) and the other located distally (SD distal to CenSat; distance > 250 kb). We applied a coverage cutoff to analyze pairs where >80% of CpG sites were covered in both copies. For each pair, DNA methylation difference was calculated as methylation of SD near CenSat minus that of SD distal to CenSat, with negative values indicating hypomethylation of SD near CenSat. Statistical significance was assessed using paired t-tests and Wilcoxon tests, with p-values and the number of SD pairs analyzed reported for each tissue. In sperm and brain, pericentromeric copies show distinct and statistically significant hypomethylation.

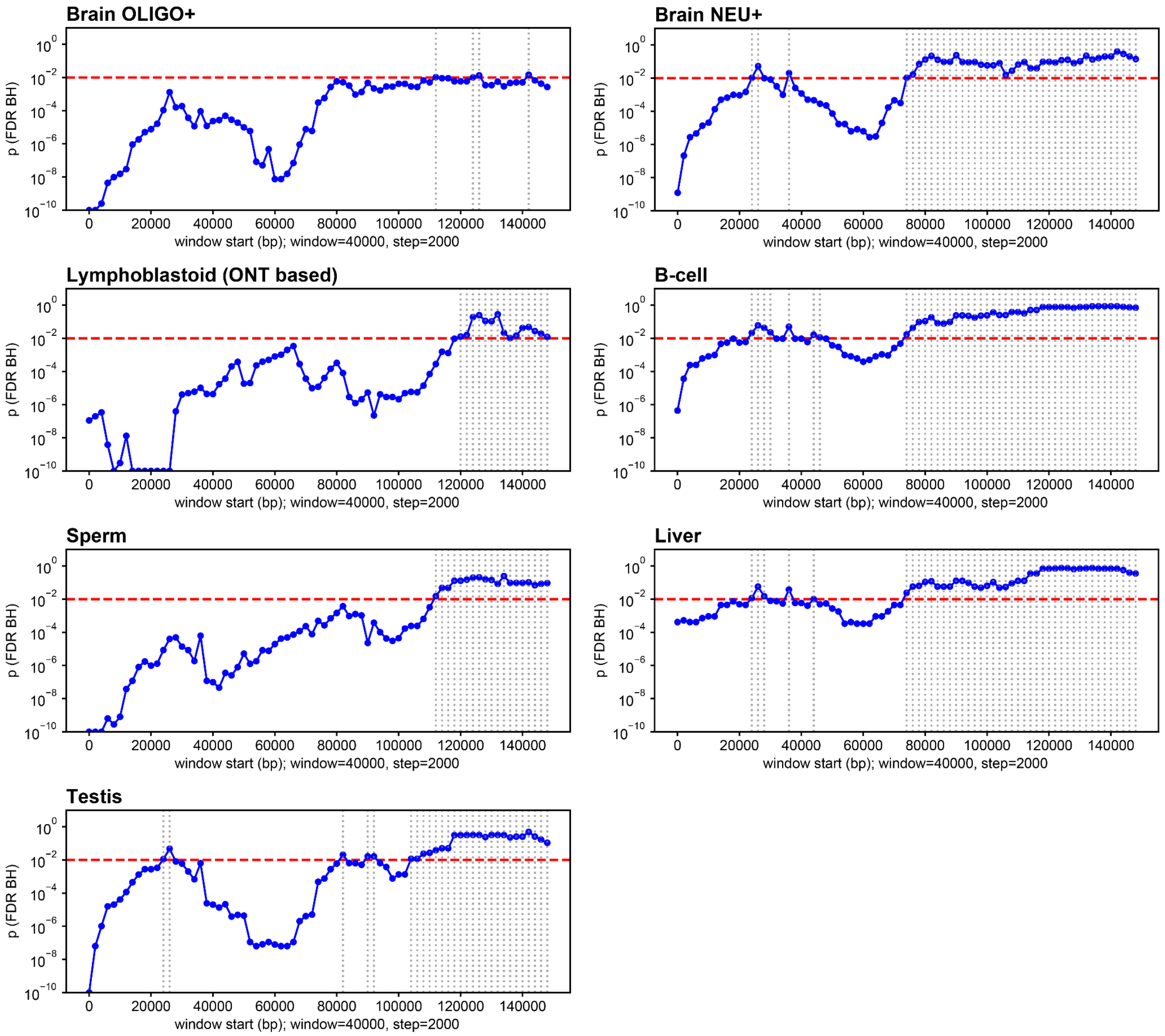


**Fig. S12. Statistical tests assessing DNA methylation divergence between SD copies in relation to the distance from satellite sequences.** Segmental duplication (SD) pairs were used to estimate the extent of peri/centromeric hypomethylation into non-satellite regions. Highly identical SD pairs (sequence identity >95%) were compared for DNA methylation in each tissue, with one copy located near satellite arrays (SD near CenSat; 0 < distance from satellite array < 100 kb) and the corresponding copy located distally (SD distal to CenSat; distance >250 kb). A coverage cutoff was applied to retain pairs in which >80% of CpG sites were covered in both copies. A sliding-window analysis across highly identical SD pairs (40-kb windows, 2-kb step size; spanning 0–150 kb from the satellite/non-satellite boundary) compared pericentromeric copies with their corresponding distal copies located ≥250 kb from satellite arrays. Statistical significance was assessed using Wilcoxon signed-rank tests with Benjamini-Hochberg FDR correction. Horizontal red dashed lines indicate an FDR-adjusted p-value threshold of 0.01, and vertical gray dashed lines indicate nonsignificant comparisons (FDR >0.01). SD copies near CenSat tended to be significantly more hypomethylated than their distal counterparts, with the magnitude of hypomethylation showing a distance-dependent pattern.

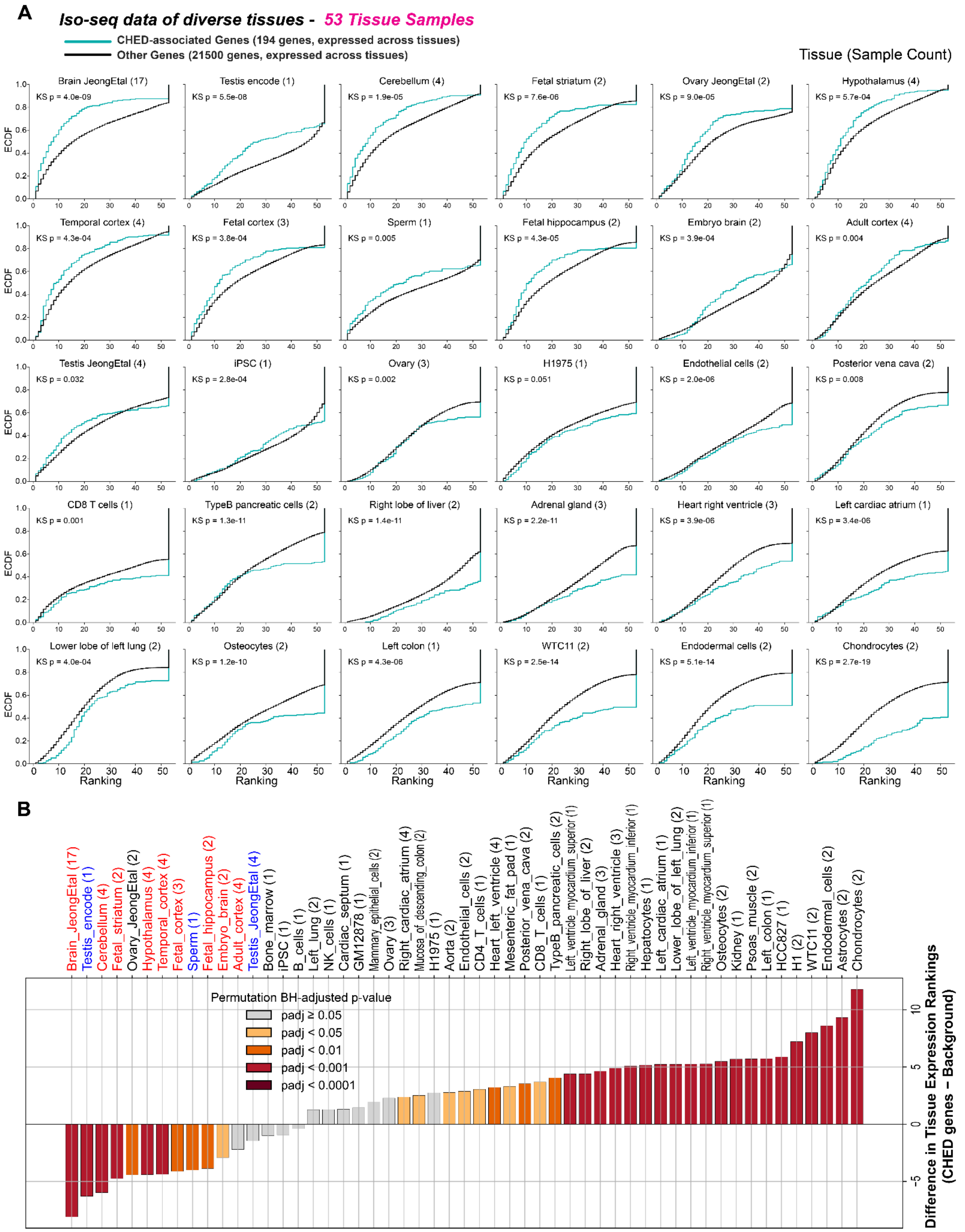


**Fig. S13. Cross-tissue expression profiles of CHED-associated genes in Iso-Seq datasets.** (A) For each tissue, relative expression across 53 long-read iso-seq tissue samples was ranked from 1 to 53 (1 = highest expression among tissues; 53 = lowest). Data are from ((*29*, *72*, *84*, *85*) and other publicly available data. Table S12). Expression-rank distributions for CHED-associated

genes and background genes elsewhere in the genome were compared using Kolmogorov-Smirnov tests of ECDFs. In testis, sperm, and multiple brain samples, CHED-associated genes showed greater enrichment than background genes, whereas most somatic tissue samples exhibited relatively low CHED-associated gene activity. Thirty representative tissue samples are shown, with P-values from Kolmogorov-Smirnov tests indicated.
(B) The difference between the mean rank of CHED-associated genes and background genes was summarized for each of the 53 tissue samples. Negative values indicate higher tissue rankings (greater tissue enrichment) for CHED-associated genes relative to background genes. Testis and sperm tissues, shown in blue, and brain-related tissues, shown in red, exhibit distinct tissue-enriched expression patterns among CHED-associated genes. Statistical significance was assessed using a two-sided permutation test with 10,000 random label permutations per tissue, followed by Benjamini-Hochberg false discovery rate correction across tissues. Corrected significance values are represented by a color gradient.

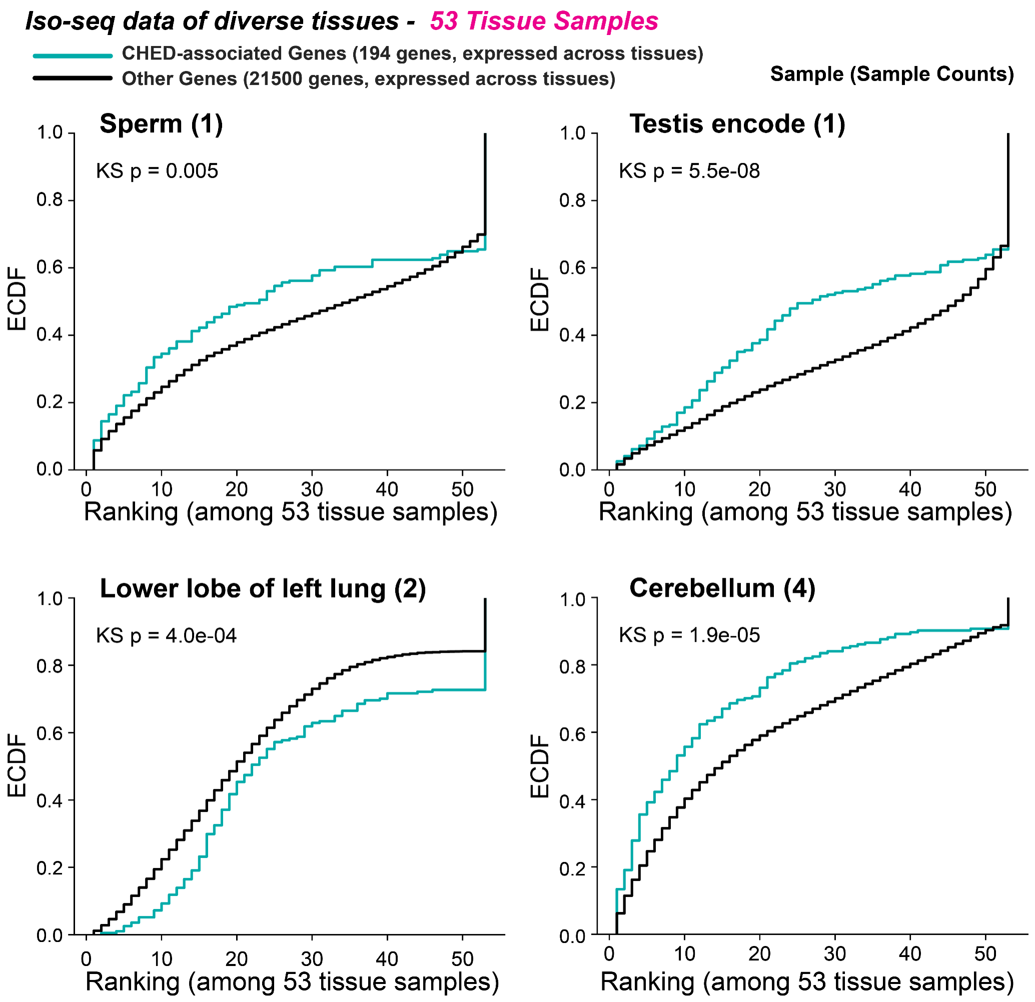


**Fig. S14. Sperm-enriched transcripts of CHED-associated genes.**
For each gene, relative expression across 53 long-read Iso-Seq tissue samples was ranked from 1 to 53, with 1 indicating the highest expression and 53 the lowest. Expression-rank distributions of CHED-associated genes and background genes elsewhere in the genome were compared using empirical cumulative distribution functions. Purified sperm Iso-Seq data showed relative enrichment of CHED-associated transcripts compared with other tissues, with similar patterns observed in brain and testis. P-values from Kolmogorov-Smirnov tests are indicated.

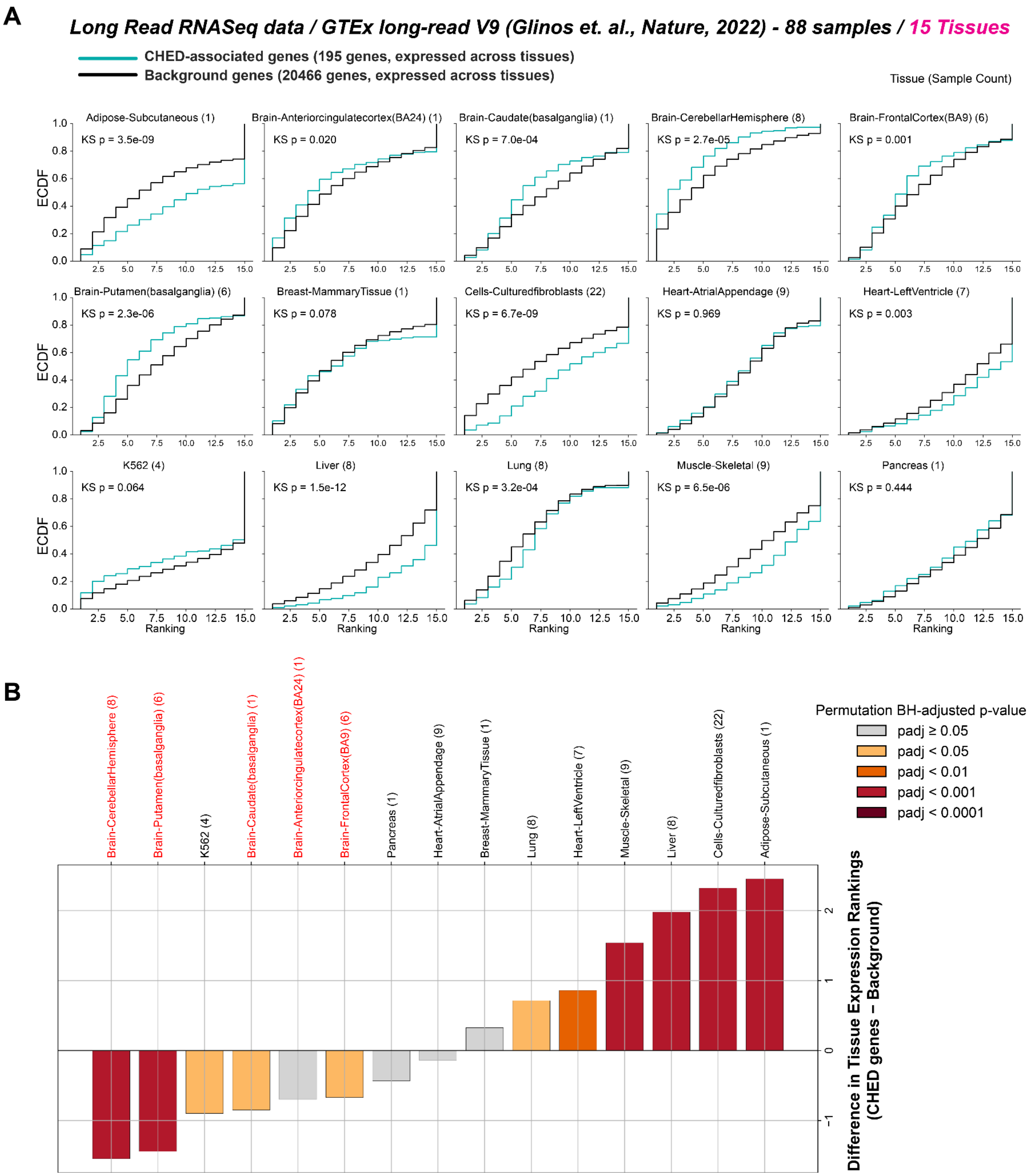


**Fig. S15. Cross-tissue expression profiles of CHED-associated genes in long-read GTEx.** (A) For each tissue, relative expression across the 15 long-read GTEx tissues was ranked from 1 to 15 (1 = highest expression among tissues; 15 = lowest). Data are from (*38*). Expression-rank distributions for CHED-associated genes and background genes elsewhere in the genome were compared using ECDFs. Across multiple brain samples, CHED-associated genes show greater tissue enrichment than background genes, whereas most other somatic tissues show relatively low CHED-associated gene activity. Sample counts are shown in parentheses. P-values from Kolmogorov-Smirnov tests are indicated.

(B) The difference between the mean rank of CHED-associated genes and background genes was summarized for each of the 15 tissues. Negative values indicate higher tissue rankings (greater tissue enrichment) for CHED-associated genes relative to background genes. Testis and brain show distinct tissue-enriched expression of CHED-associated genes. Statistical significance was assessed using a two-sided permutation test with 10,000 random label permutations per tissue, followed by Benjamini-Hochberg false discovery rate correction across tissues. Corrected significance values are represented by a color gradient.

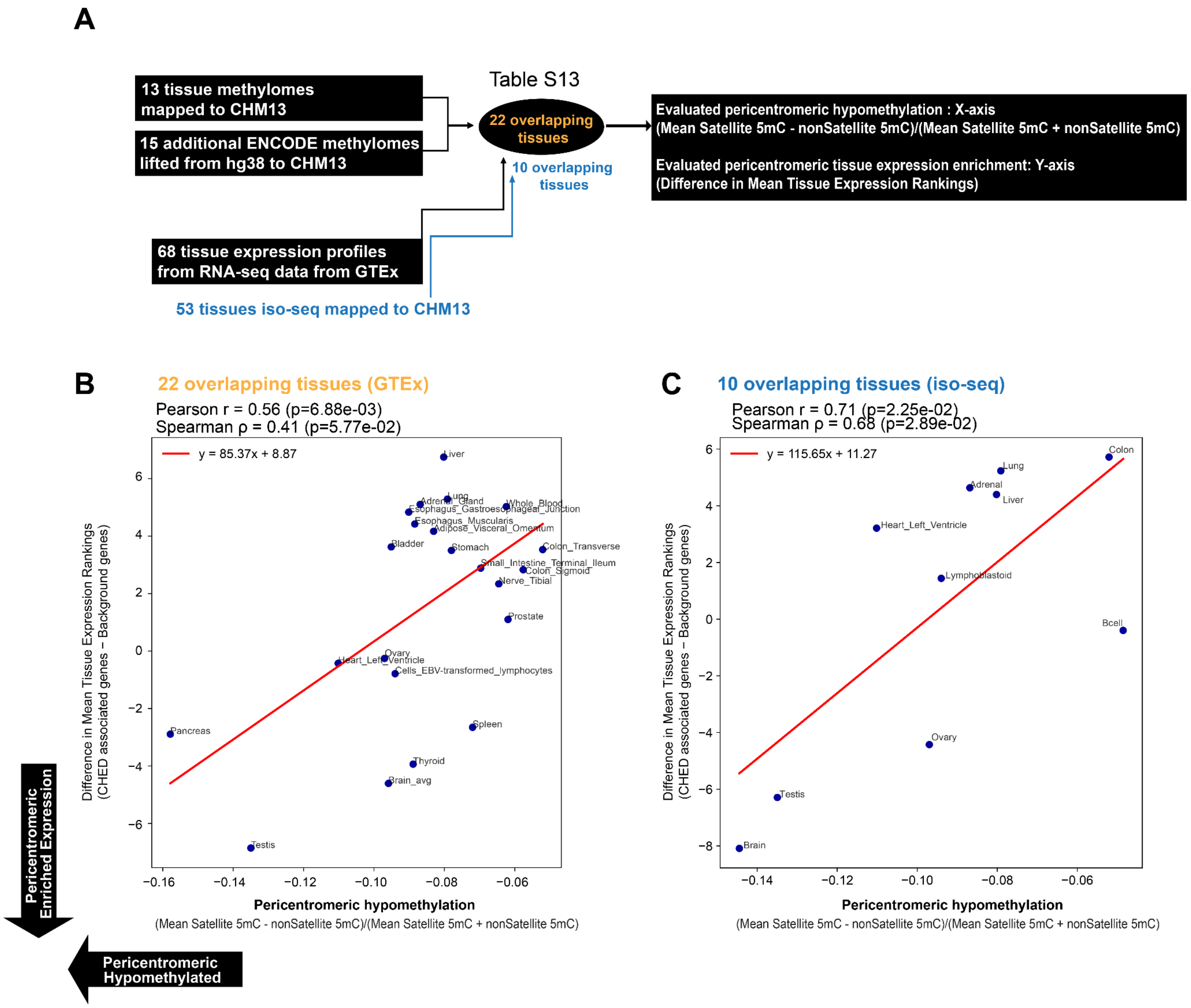


**Fig. S16. Correlation between peri/centromeric hypomethylation and tissue enrichment of CHED-associated genes.**

(A) A comprehensive set of tissue-matched methylation-expression profiles was generated for tissues with both WGBS methylation and expression data available. This included 22 tissues shared between GTEx RNA-seq datasets and methylomes, and 10 tissues shared between Iso-Seq datasets and methylomes. For each tissue, both the degree of peri/centromeric hypomethylation and the degree of tissue enrichment of CHED-associated gene expression were evaluated.

(B) For the 22 shared GTEx RNA-seq and methylome tissues, we tested the association between peri/centromeric hypomethylation and enrichment of CHED-associated gene expression. Tissue enrichment of CHED-associated gene expression was significantly correlated with the degree of peri/centromeric hypomethylation: tissues with stronger peri/centromeric hypomethylation tended to show stronger tissue-enriched expression of CHED-associated genes, as assessed by Spearman and Pearson correlations.

(C) The same analysis was performed for the 10 tissues shared between the Iso-Seq and methylome datasets. Correlation coefficients and P-values are indicated.

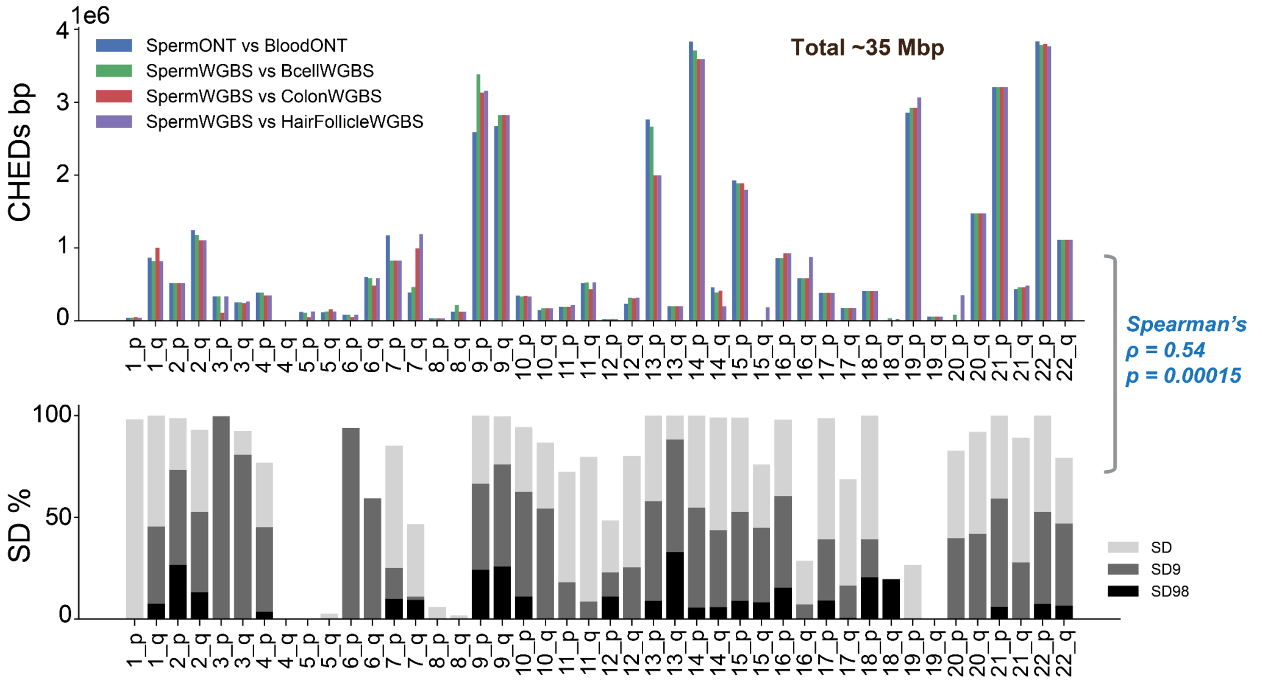


**Fig. S17. The extent of CHEDs was positively correlated with the abundance of SDs.** (Top) Base pairs of CHEDs detected on each chromosome arm in pairwise comparisons of sperm data with somatic tissues lacking extended hypomethylation: blood, B cells, colon, and hair follicle. Hypomethylated domains were highly concordant across ONT- and WGBS-based pairwise comparisons. Approximately 35 Mb of significant regions were shared across pairwise comparisons. (Bottom) Segmental duplication content calculated within 0 to 200 kb regions from satellites on each chromosome for SDs, highly identical SDs, SD95, and SD98. The amount of CHEDs was significantly associated with SD content, as determined by Spearman correlation across chromosomes.

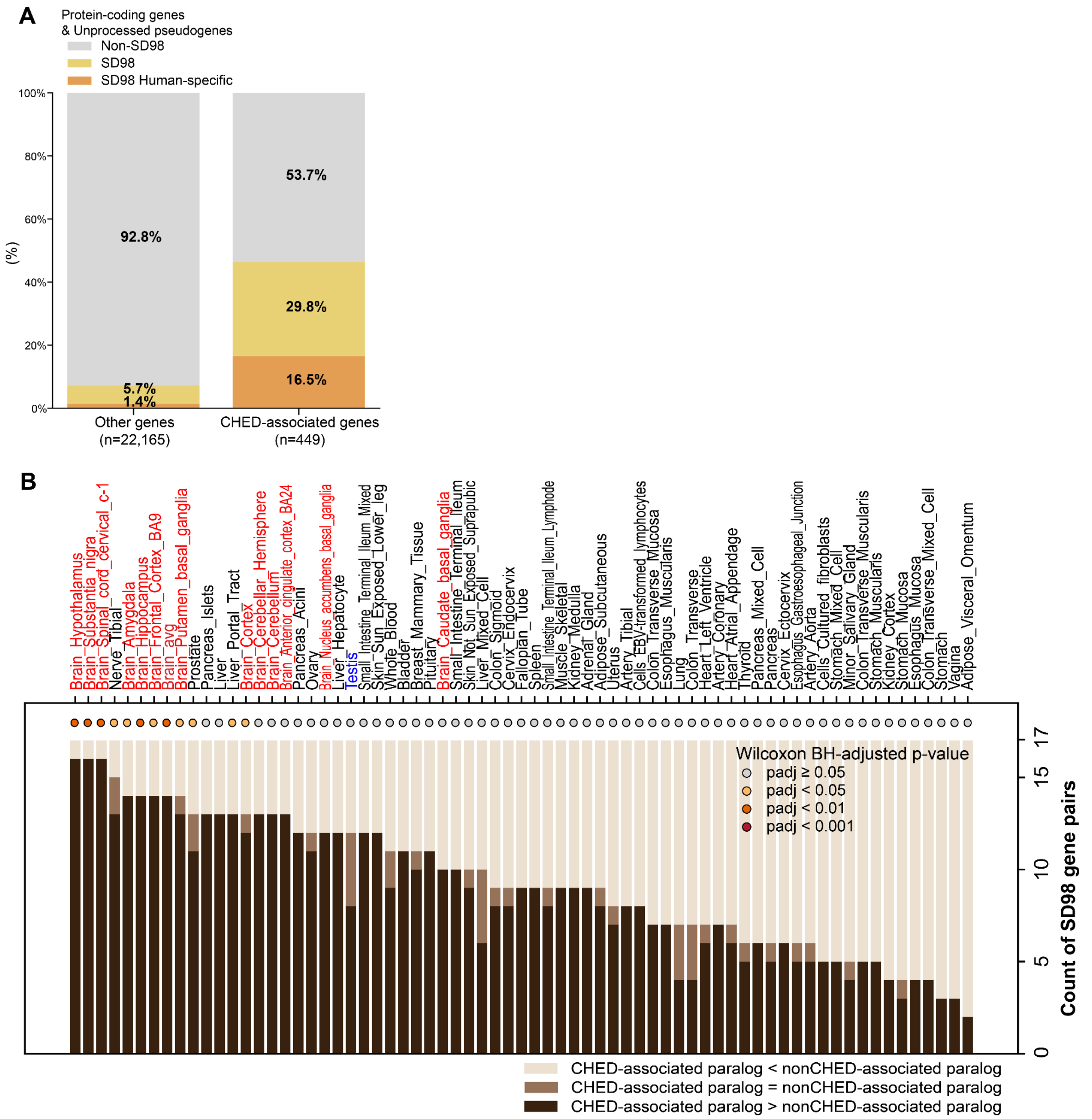


**Fig. S18. Recently duplicated genes overrepresented within CHEDs.** (A) The proportion of recently duplicated genes, defined as SD98 genes (*36*) (1,793 protein-coding genes and pseudogenes), was calculated for CHED-associated genes and other genes. Among the 449 CHED-associated protein-coding genes and pseudogenes, recently duplicated genes (SD98) were highly enriched, and many were also human-specific based on synteny with chimpanzees. (B) SD98 gene pairs in which one paralog is CHED-associated and the other is nonCHED-associated were compiled, yielding 84 SD98 paralog pairs in total. Of these, 17 had available cross-tissue expression values for both paralogs. Tissue-enrichment divergence for these 17 pairs was quantified using tissue expression ranks by counting the number of pairs in which the CHED-associated paralog had a higher tissue rank than its nonCHED-associated counterpart. Testis is shown in blue, and brain tissues are shown in red. Statistical significance was assessed using a one-sided Wilcoxon signed-rank test comparing whether expression ranks were higher for CHED-associated paralogs than for nonCHED-associated paralogs. FDR-corrected significance is indicated by colored circles.

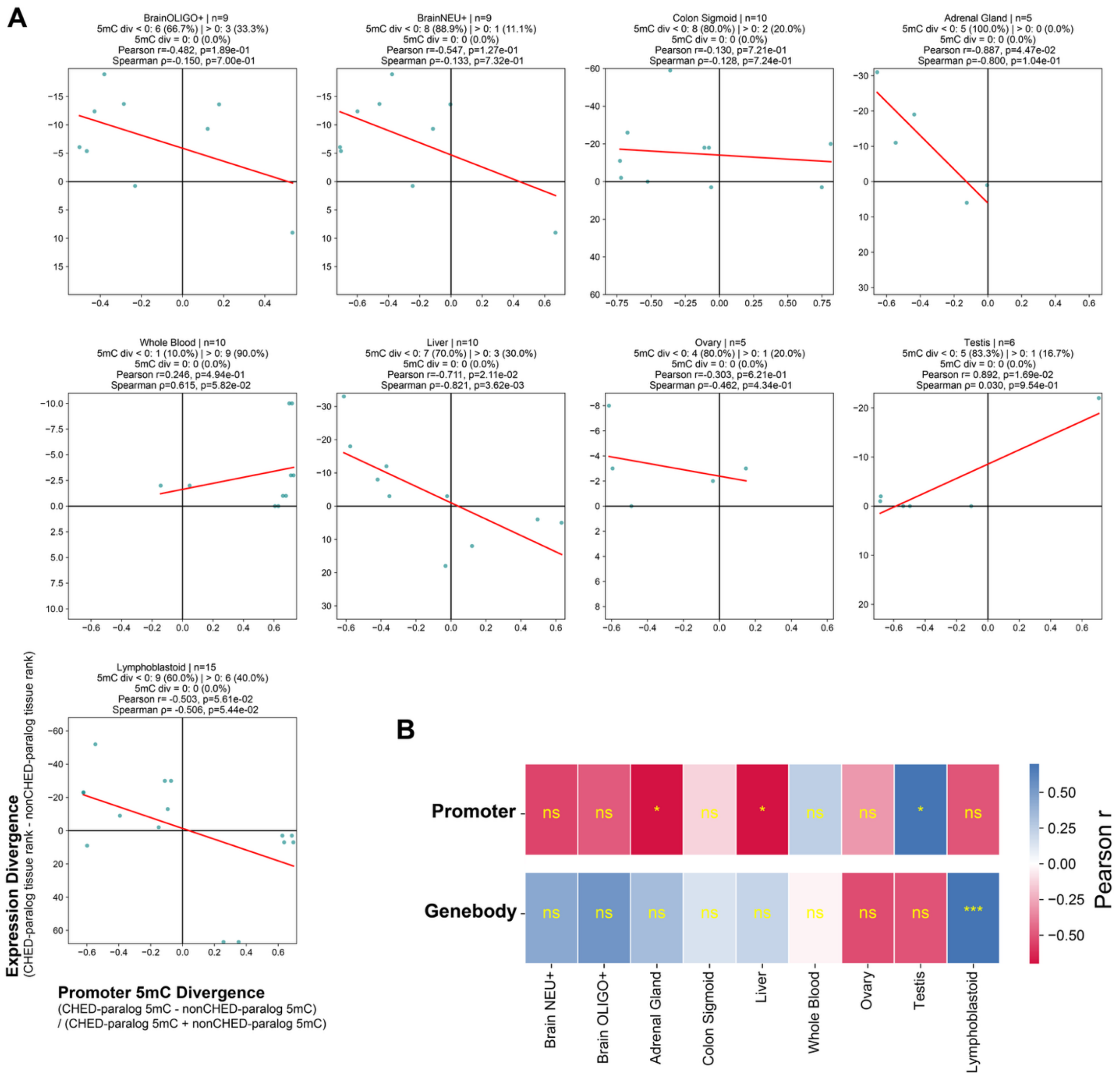


**Fig. S19. Correlation between DNA methylation and expression divergence in SD98 gene pairs across matched tissues.**

(A) SD98 gene pairs were compiled in which one paralog was pericentromeric (CHED-associated) and the other was non-pericentromeric. In each tissue, we analyzed highly identical SD98 paralog pairs for which both copies were expressed and DNA methylation data were available for both copies, with >66% of CpG sites covered in each copy. Distributions of expression divergence and promoter DNA methylation divergence are shown for covered SD98 pairs across nine tissues with both DNA methylome and GTEx expression data available. Negative x-axis values indicate pairs in which the CHED-associated copy is hypomethylated, and negative y-axis values indicate pairs in which the CHED-associated copy has a higher expression rank than the nonCHED-associated paralog. The y-axis is inverted so that higher expression ranks are displayed upward. At the top of each plot, the tissue, number of available data points, number and percentage of SD98 pairs in which the CHED-associated paralog is hypomethylated relative to its non-CHED counterpart, and the results of Pearson and Spearman correlation tests are shown. Many CHED-associated paralogs show greater promoter hypomethylation than their non-CHED-associated counterparts, and this hypomethylation is associated with higher tissue-enriched expression, resulting in an overall negative correlation across many tissues.

(B) The same analysis was performed and summarized for both promoter and gene-body DNA methylation across tissues. The association between DNA methylation divergence and expression divergence was summarized using Pearson correlation coefficients across tissues. A negative correlation indicates that hypomethylation of the CHED-associated paralog is associated with higher tissue activity. Promoter methylation divergence tended to be negatively correlated with expression divergence between CHED-associated and non-CHED-associated paralogs across many tissues.

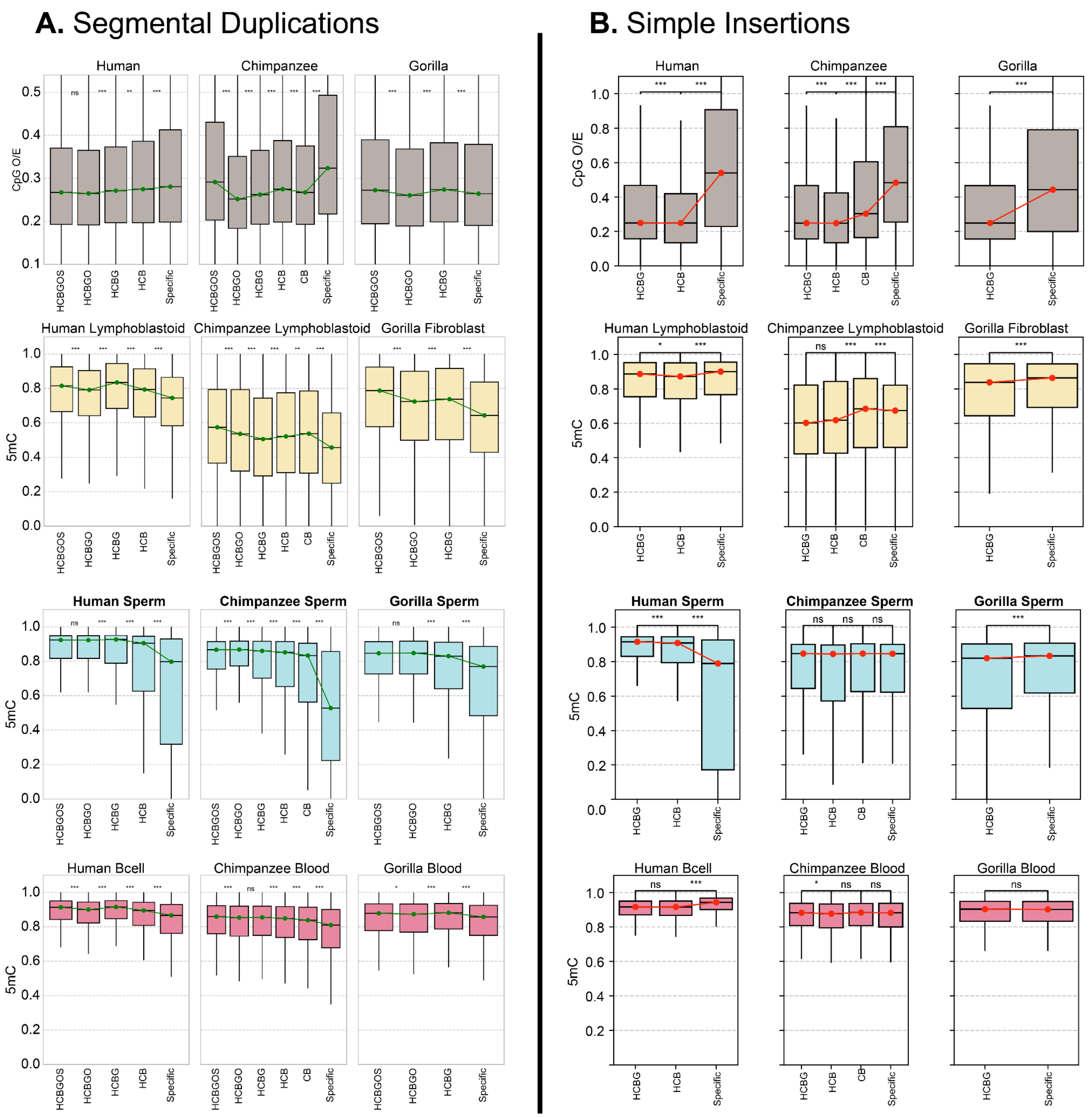


**Fig. S20. Germline hypomethylation in evolutionarily recent SDs and insertions.** For each evolutionary node, shared and lineage-specific elements were identified for segmental duplications (A) and simple insertions (B). Older elements are positioned further to the left on the x-axis. CpG O/E (gray) and DNA methylation levels in somatic cell lines (yellow), sperm (blue), and B cells/blood (red) are compared, revealing a gradual decrease in CpG O/E and pronounced sperm hypomethylation in lineage-specific SDs, patterns not observed in B-cell/blood samples. A similar pattern of high CpG O/E and sperm hypomethylation is observed in evolutionarily recent simple insertions, particularly in humans.

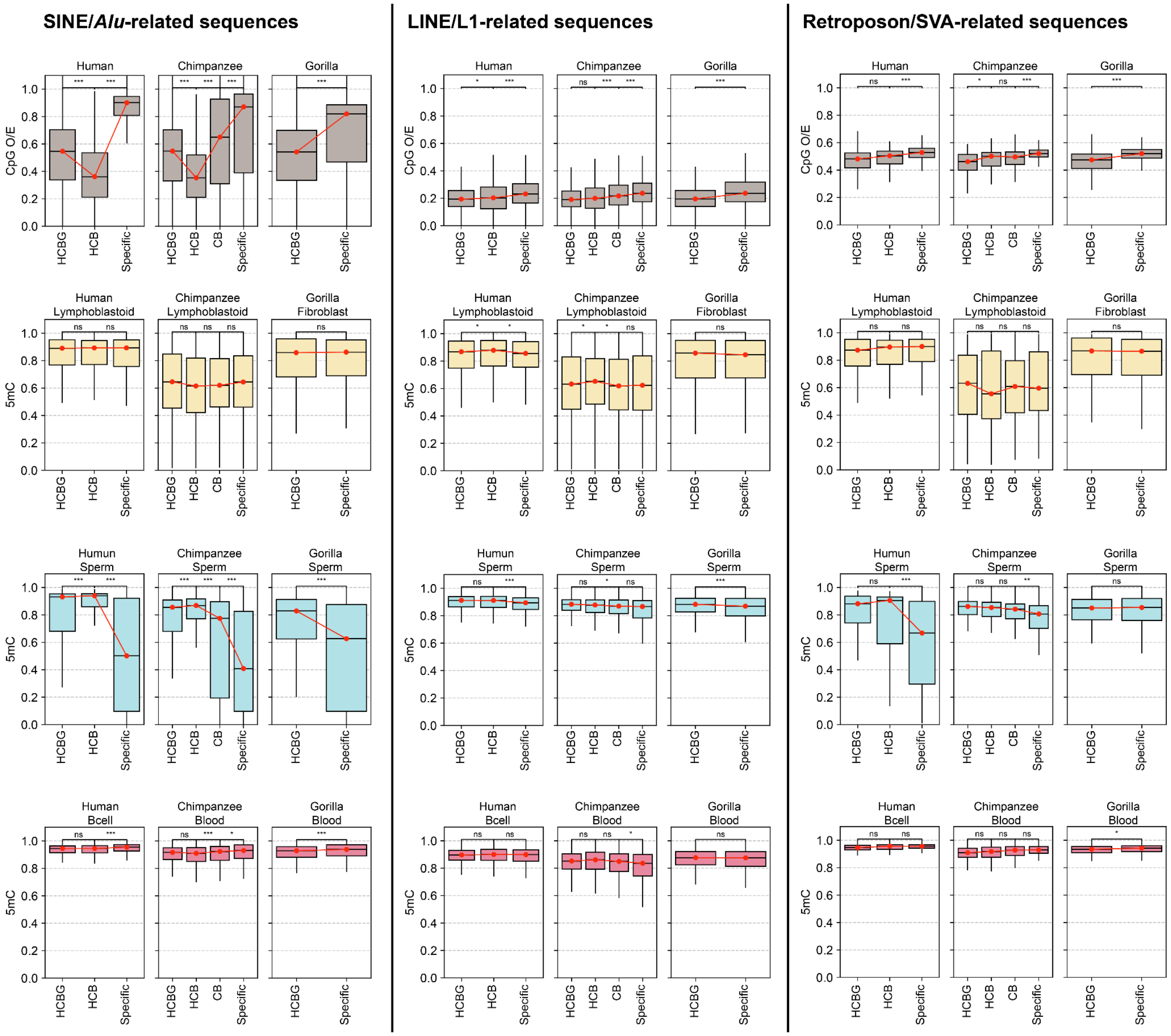


**Fig. S21. Germline hypomethylation in evolutionarily recent insertions across different sequence classes.** Given the high sequence heterogeneity of simple insertions, we further characterized and compared recurrent sequence classes within them based on repeat composition, focusing on active transposable element classes including SINE/Alu, LINE/L1, and retroposon/SVA. Repeat elements fully contained within simple insertion intervals were identified and classified by repeat type. CpG O/E (gray) and DNA methylation levels in somatic cell lines (yellow), sperm (blue), and B cells/blood (red) are compared across classes. All classes show higher CpG O/E in lineage-specific elements together with distinct sperm hypomethylation, patterns that are not observed in somatic samples.

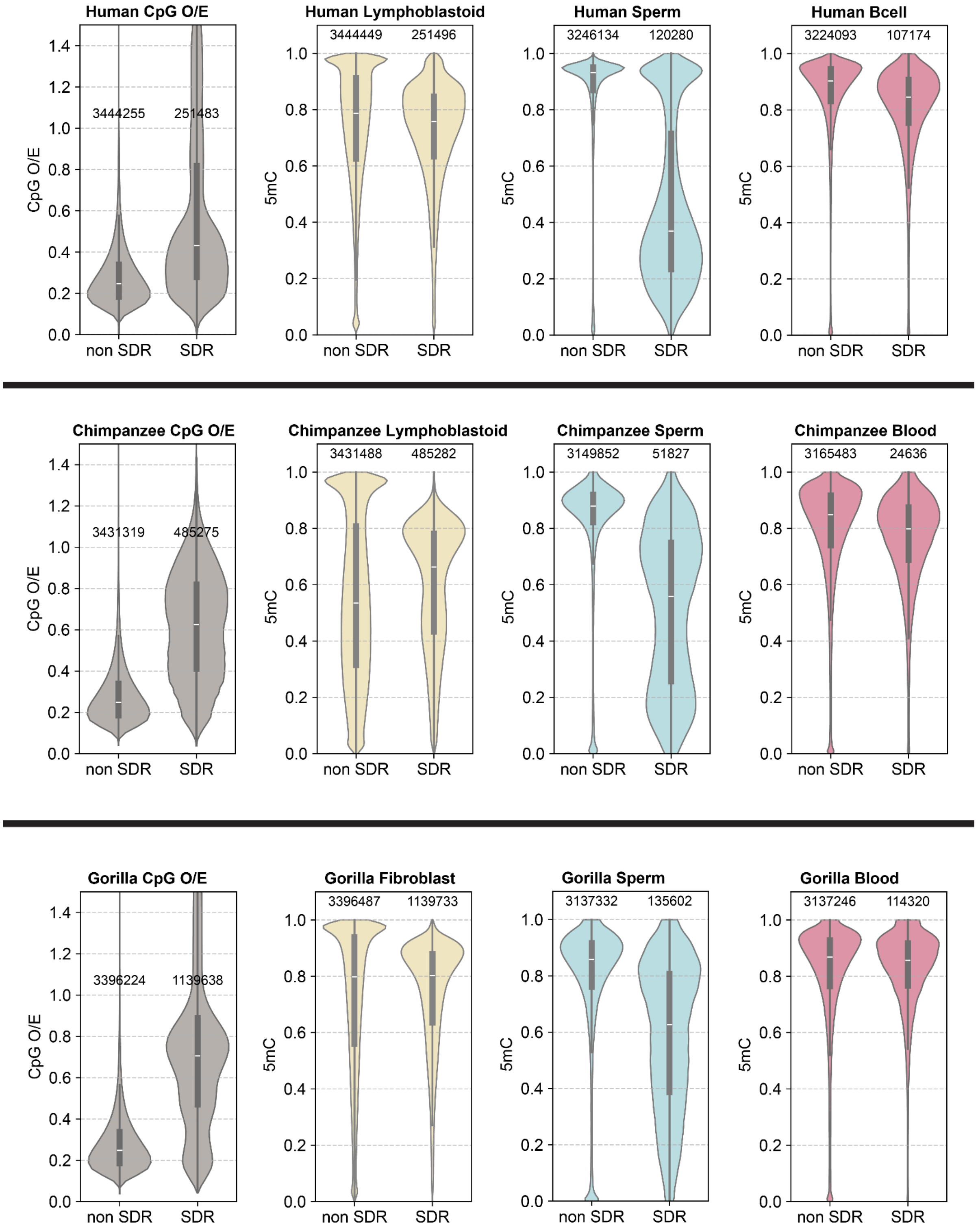


**Fig. S22. Germline hypomethylation in structurally divergent regions (SDRs).**
CpG O/E and DNA methylation were compared between non-SDRs and SDRs across tissues. CpG O/E (gray) and DNA methylation levels in somatic cell lines (yellow), sperm (blue), and B cells/blood (red) are shown. SDRs show less CpG depletion and lower sperm methylation, a pattern not observed in lymphoblastoid cells. Analyses were performed in human, chimpanzee, and gorilla.

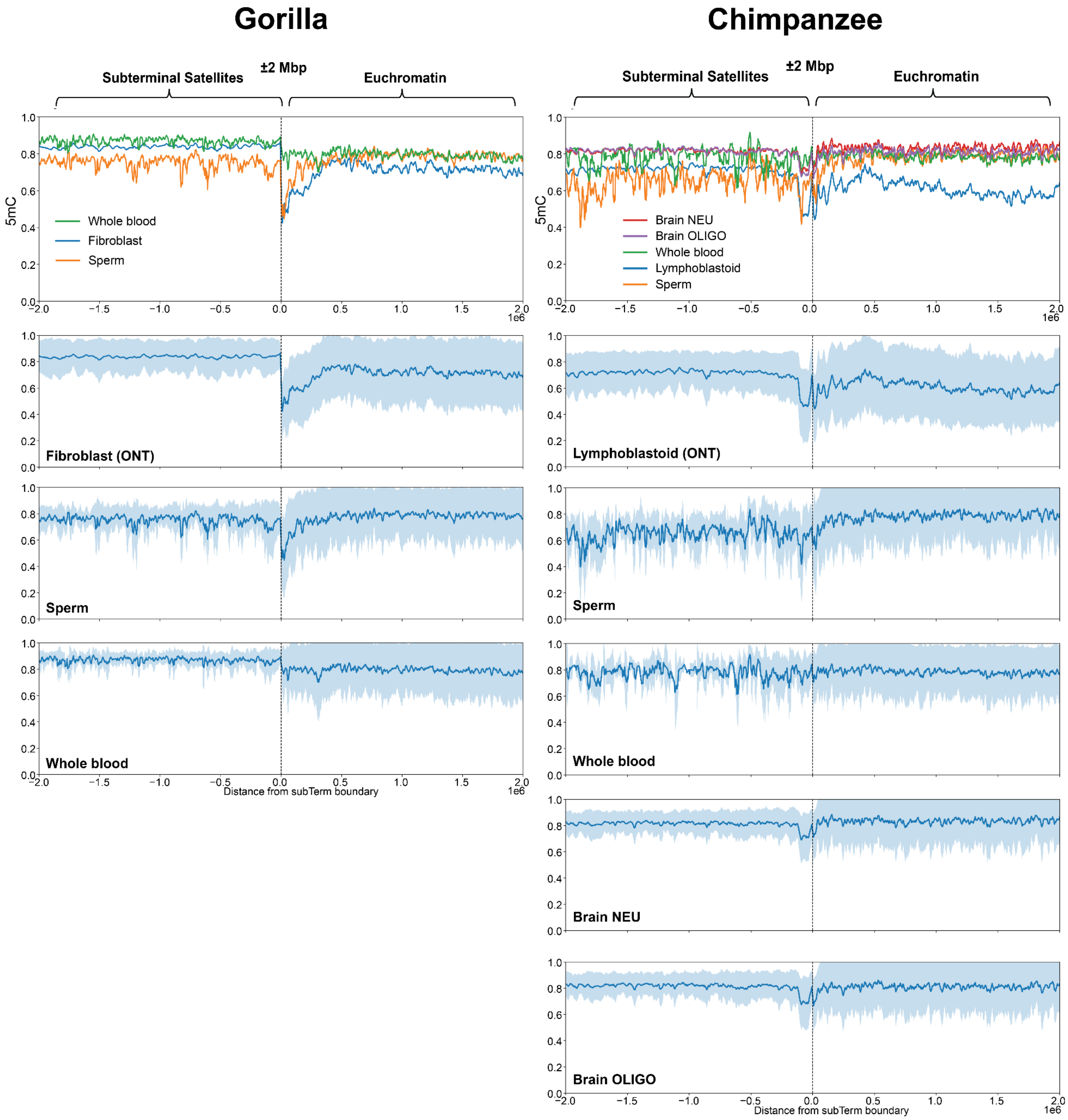


**Fig. S23. Methylation profiles within 2 Mb of the euchromatin-heterochromatin transition for chromosomes with subterminal caps in gorilla (left) and chimpanzee (right).** The euchromatin-heterochromatin boundary is defined here as the last proximal subterminal pCht satellite unit (*35*), and the surrounding 2 Mbp regions were investigated. For gorilla, methylation landscapes for whole-blood, fibroblast, and sperm samples are compared together (top) and plotted separately below. For chimpanzee, lymphoblastoid, sperm, whole-blood, brain NEU+, and brain OLIGO+ samples are compared. In gorilla, an immediate and sharp drop in methylation at the boundary is observed in fibroblast and sperm samples, but not in the whole-blood sample. Similarly, in chimpanzee, hypomethylation near the boundary shows variable patterns across tissues.
